# Fetal context conveys heritable protection against MLL-rearranged leukemia that depends on MLL3

**DOI:** 10.1101/2025.03.11.642680

**Authors:** Jonny Mendoza-Castrejon, Wei Yang, Elisabeth Denby, Helen Wang, Emily B. Casey, Rohini Muthukumar, Riddhi M. Patel, Jihye Yoon, Yanan Li, J. Michael White, Ran Chen, Luis F.Z. Batista, Jeffrey A. Magee

## Abstract

*MLL* rearrangements (*MLL*r) are the most common cause of congenital and infant leukemias. *MLL*r arise prior to birth and require few cooperating mutations for transformation, yet congenital leukemias are 10-fold less common than infant leukemias and >100-fold less common than childhood leukemias overall. This raises the question of whether mechanisms exist to suppress leukemic transformation during fetal life, thereby protecting the developing fetus from malignancy during a period of rapid hematopoietic progenitor expansion. Here, we use mouse models to show that fetal MLL::ENL exposure creates a heritable, leukemia-resistant state. MLL::ENL imposes a negative selective pressure on fetal hematopoietic progenitors. It leads to postnatal loss of self-renewal gene expression and enhanced myeloid differentiation that precludes transformation. These changes do not occur when MLL::ENL is induced shortly after birth, and transformation proceeds efficiently in this context. The fetal barrier to transformation is enforced by the histone methyltransferase MLL3. It can be overcome by cooperating mutations, such as *Nras^G12D^*, or through somatic or germline inactivation of MLL3. Heritable fetal protection against leukemic transformation may explain the low incidence of congenital leukemias in humans despite prenatal *MLL* rearrangement.

## INTRODUCTION

Infant leukemias are rare, difficult-to-treat malignancies that occur early in life and are genetically distinct from later childhood and adult leukemias (1,2). Approximately 50% of infant acute myeloid leukemias (AML) and ∼70% of infant B-cell acute lymphoblastic leukemias (B-ALL) harbor an *MLL/KMT2A* rearrangement (*MLL*r) as the primary driver mutation (3–5). *MLL*r arise as translocation mutations that fuse the N-terminus of the MLL protein to one of several potential C-terminal partners (6). The resulting fusion oncoproteins ectopically activate *HOX* family genes, *MEIS1* and other self-renewal genes in either myeloid or B-cell progenitors to drive leukemogenesis (7–9). Aside from *MLL*r, the genomes of infant leukemias typically carry few additional cooperating mutations (5,10,11). This contrasts with the higher mutation burdens of adolescent and adult leukemias. It suggests that fetal and neonatal progenitors require only a few mutations to transform into malignant cells, once an *MLL*r has been established.

Several lines of evidence implicate a fetal cell of origin for *MLL*r infant leukemias. Retrospective analyses of newborn blood screens showed that infant B-ALL patients had molecular evidence of the *MLL*r at birth, prior to the onset of frank leukemia (12–15). Furthermore, monozygotic twin studies showed that when one twin develops infant leukemia, the second twin also develops infant leukemia with near 100% concordance (16,17). Together, these data show that *MLL*r occur before birth and raise the question of whether fetal/neonatal progenitors might be particularly prone to transformation. Indeed, prior studies with either mouse or human models have shown that MLL fusion proteins, including MLL::ENL, MLL::AF9 and MLL::AF4, all transform fetal/neonatal hematopoietic progenitors more efficiently than adult progenitors (18–20). This has become a working explanation for both the high frequencies at which *MLL*r occur in infant leukemias and the relatively small number of cooperating mutations required for transformation.

Despite its simplicity, other lines of evidence argue against a model in which fetal blood progenitors are efficiently transformed by *MLL*r mutations. While *MLL*r cause a high percentage of infant leukemias, the overall incidence of *MLL*r infant leukemia is actually quite low. The incidence of infant AML is only ∼15 cases per million, approximately 20-fold lower than the incidence of AML in a 70-year-old (21). The incidence of congenital leukemia is lower still (∼3 cases per million) (21,22). When also accounting for lymphoblastic leukemias, the incidence of congenital leukemias is 10-fold lower than infant leukemias and >100-fold lower than childhood leukemia overall (21). This is counterintuitive for two reasons. First, fetal hematopoietic stem cells (HSCs) and multipotent progenitors (MPPs) divide more rapidly than postnatal HSCs and MPPs, thus affording *MLL*r clones ample opportunity to expand prior to birth. Second, as noted above, only a small number of cooperating mutations are required for infant leukemias to transform. In principle, these factors should cause congenital leukemias to occur more often than childhood or adult leukemias, rather than infrequently. It is possible that the low incidence of congenital leukemias simply reflects high integrity of the genome early in life, despite high rates of cell proliferation, but the observation nevertheless raises the question of whether additional mechanisms exist to suppress, rather than potentiate, leukemic transformation during fetal ontogeny.

In prior work, we used a Doxycycline (DOX)-inducible model of MLL::ENL-driven AML to test whether AML initiation efficiency changes with age (20). We found that mice with a tetracycline-regulated MLL::ENL transgene (Tet-Off-ME mice) developed AML with greatest penetrance when MLL::ENL was first induced shortly after birth, rather than before birth (20). This observation suggests that fetal progenitors may resist AML initiation. The underlying reason for this discrepancy remains unclear. We and others showed that a fetal-specific RNA binding protein, LIN28B, can actively suppress MLL::ENL-driven leukemogenesis (20,23). However, in subsequent loss-of-function studies, we showed that fetal progenitors are protected from leukemic transformation even in the absence of LIN28B (24). Other mechanisms must convey protection against transformation when MLL::ENL is expressed before birth.

We now show that MLL::ENL imposes a negative selective pressure on fetal progenitors that then conveys a heritable, AML-resistant state. MLL::ENL suppresses fetal HSC/MPP proliferation to an extent not seen after birth. Furthermore, fetal MLL::ENL induction leads to a postnatal decline in expression of select stemness genes (e.g., *Pbx1*, *Nkx2-3, Myct1* and *Hlf*), and it enhances postnatal myeloid priming. Interestingly, the enhanced myeloid priming reflects MLL::ENL-driven changes in gene expression but not chromatin accessibility. In other words, the gene expression profiles of myeloid progenitors – at every stage from hematopoietic stem cell (HSC) to granulocyte-monocyte progenitor (GMP) – reflect a more differentiated state than would be inferred based on chromatin accessibility. The heritable, AML-resistant state is enforced by MLL3, an epigenetic regulator that promotes myeloid differentiation. Even monoallelic *Mll3* deletions are sufficient to restore efficient transformation after fetal MLL::ENL induction.

Our data argue for a nuanced interpretation of how ontogeny influences leukemogenesis. We show that fetal developmental context imposes a barrier to transformation that may limit the incidence of congenital and infant leukemias, but the barrier can be overcome by cooperating mutations or loss of key epigenetic regulators. This model of fetal-specific cancer suppression may help explain why developing organ systems, such as the fetal hematopoietic system, can undergo exponential expansion during ontogeny with only minimal risk for neoplastic transformation.

## RESULTS

### MLL::ENL induction in fetal progenitors conveys heritable protection against transformation

We previously showed, using a Tet-Off model for MLL::ENL induction, that postnatal MLL::ENL induction led to faster and more penetrant AML initiation than fetal induction (20). This raised the question of whether cooperating mutations, such as the RAS pathway mutations that frequently co-occur in *MLL*r infant leukemias, could mitigate the difference between fetal and postnatal transformation efficiency. Given that DOX can take time to clear in a Tet-Off system, we transitioned to a Tet-On system for these experiments to enable more precise control of MLL::ENL induction. We generated *Vav1-Cre*; *Rosa26^LSL-rtTA-IRES-mKate2^*; *Col1a1^TetO-MLL::ENL^* (Tet-On-ME) and *Vav1-Cre*; *Rosa26^LSL-rtTA-IRES-mKate2^*; *Col1a1^TetO-MLL::ENL^*; *Nras^LSL-G12D^* (Tet-On-ME/Nras^G12D^) mice so that we could induce MLL::ENL expression in fetal or neonatal hematopoietic progenitors by administering DOX. We induced MLL::ENL expression at either embryonic day (E)10.5 or postnatal day (P)0 by feeding pregnant/nursing mothers DOX chow, and we monitored survival of the progeny. For Tet-On-ME mice, MLL::ENL induction immediately after birth caused fully penetrant AML with a median survival of ∼3 months, whereas MLL::ENL induction before birth led to incompletely penetrant AML initiation with a median survival of ∼5 months (Fig. 1A). These survival curves were similar to what we previously observed using the Tet-Off model (20). For Tet-On-ME/Nras^G12D^ mice, AML developed with complete penetrance after both fetal and postnatal MLL::ENL induction (Fig. 1B). However, Tet-On-ME/Nras^G12D^ mice survived longer after fetal MLL::ENL induction, indicating that fetal developmental context can suppress AML initiation even in the presence of a cooperating mutation.

**Figure 1.**
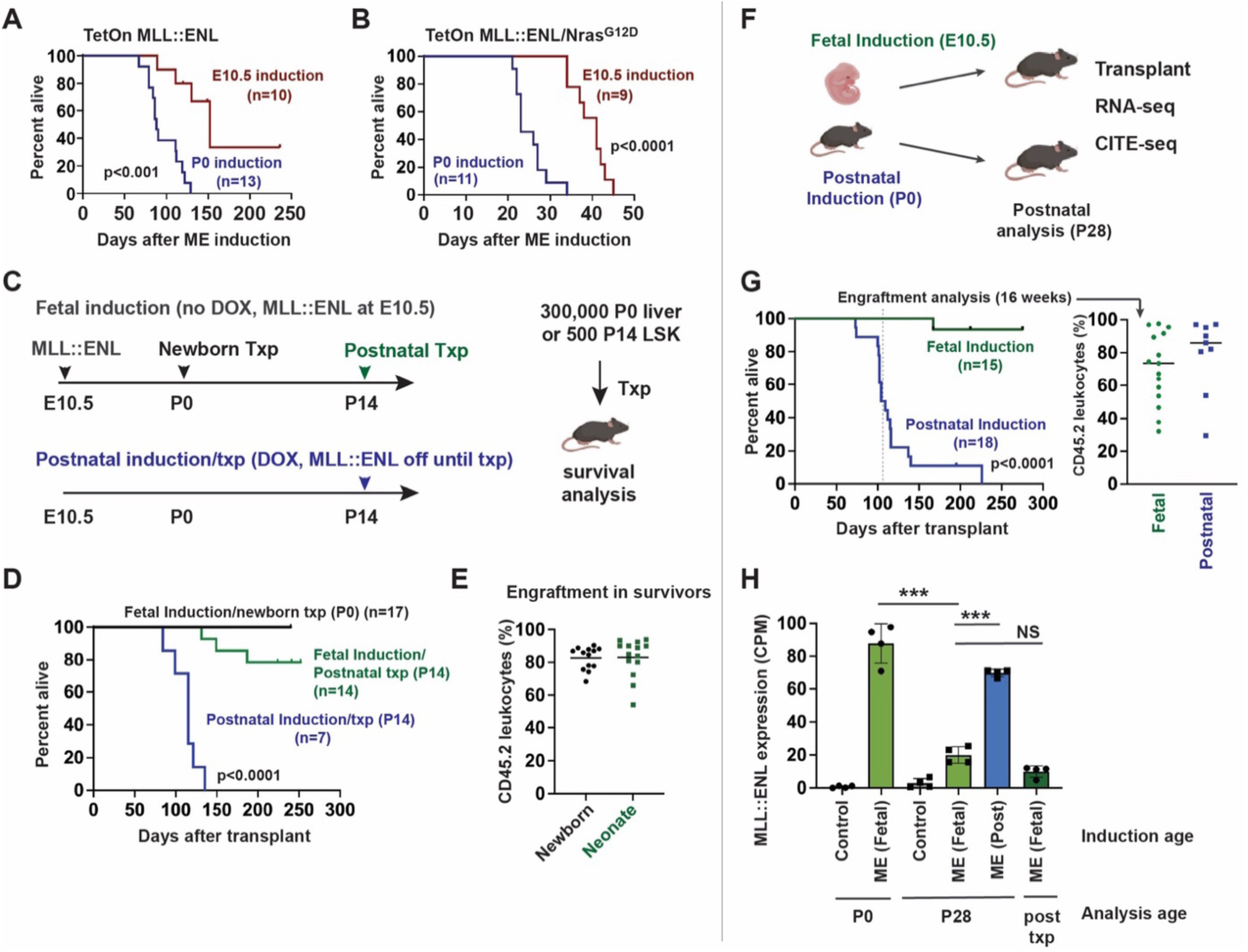
Fetal MLL::ENL induction conveys heritable protection against leukemogenesis. (A, B) Kaplan-Meier survival curves after E10.5 or P0 MLL::ENL induction using Tet-On-ME and Tet-On-ME/Nras^G12D^ models. (C) Summary of experimental set-up for MLL::ENL induction and transplantation experiments. (D) Kaplan-Meier survival curves for recipient mice transplanted with 300,000 P0 liver cells or 500 LSK cells after MLL::ENL induction at E10.5 or P14. (E) Donor (CD45.2) leukocyte engraftment in surviving recipient mice at 8 months after transplantation. (F) Summary of experimental set-up for panels G and H, and for CITE-seq experiments in Fig. 2. (G) Kaplan-Meier survival curves and 16-week donor chimerism for recipient mice after MLL::ENL induction at E10.5 or P0 and transplantation at P28. (H) MLL::ENL transgene expression (counts per million; CPM) based on RNA-seq. n= 4 biological replicates per group, ***p<0.001 by one-way ANOVA with Tukey’s posthoc test. For all survival curves, group sizes and p-values are shown in the panels. p-values were calculated by the log rank test.

The survival data raised the question of whether MLL::ENL elicits changes in fetal progenitors that persist and convey long-lasting, heritable resistance to AML. To test this possibility, we transitioned back to the Tet-Off-ME model to simplify the DOX treatment regimen. We induced MLL::ENL at E10.5 and then transplanted either P0 or P14 progenitors into lethally irradiated adult recipient mice (Fig. 1C, Supp. Fig. 1A). None of the recipients of P0 liver cells, and only a minority of recipients of P14 Lineage^-^ Sca1^+^Kit^+^ (LSK) cells, developed AML despite high levels of donor cell engraftment (Fig. 1D, E, Supp. Fig. 1B). For comparison, we also induced MLL::ENL at P14 in conjunction with the transplant. In this case, all recipient mice developed AML (Fig. 1D). We performed similar experiments with the Tet-On-ME/Nras^G12D^ model. We induced MLL::ENL by administering DOX chow at E10.5 or P21 and then transplanted progenitors into DOX-fed recipients at P0 or P28, respectively. The cooperating *Nras^G12D^* mutation did enable AML formation in all recipients, but the latency was again longer after fetal MLL::ENL induction (Supp. Fig. S1C). Thus, fetal MLL::ENL induction elicits heritable changes in hematopoietic progenitors that limit transformation potential. The barrier to transformation persists after birth and even after transplantation into adult recipients.

Age-related changes in MLL::ENL transgene expression could contribute to differences in AML penetrance. To assess MLL::ENL expression, we repeated the fetal and postnatal induction and transplantation experiments. We induced MLL::ENL before or after birth, as in the prior experiment, and then isolated LSK cells at P28 for transplantation and RNA-sequencing (RNA-seq) (Fig. 1F). We also isolated LSK cells from P0 mice and from recipient mice (∼8 months post-transplant) for RNA-seq to quantify expression of the human MLL::ENL transgene. Transplantation assays redemonstrated a profound difference in survival after fetal and postnatal MLL::ENL induction, even though the donor mice in this case were identically aged (Fig. 1G). After fetal induction, MLL::ENL was highly expressed at P0, but expression levels dropped by P28 and were significantly lower than was observed after postnatal induction (Fig. 1H). MLL::ENL expression remained low after fetal induction even in transplant recipients (Fig. 1H). Postnatal and post-transplant declines in transgene expression were not observed when another transgene – Tet-Off-H2B-GFP – was expressed from the *Col1a1* locus (Supp. Fig. 2). Thus, age-related differences in MLL::ENL expression are not attributable to non-specific differences in regulation of the *Rosa26^LSL-tTA^* or *Col1a1* loci. The data suggest that MLL::ENL imposes a negative selective pressure on fetal progenitors that favors outgrowth of cells with low transgene expression and limited transformation potential.

### Fetal MLL::ENL induction causes precocious myeloid differentiation and loss of stemness gene expression after birth

To understand how MLL::ENL differentially regulates gene expression and cell identity after fetal and postnatal induction, we performed Cellular Indexing of Transcriptomes and Epitopes followed by sequencing (CITE-seq) on P28 Lineage^-^c-Kit^+^ (LK) progenitors after fetal or postnatal MLL::ENL induction (Fig. 1F). The data were analyzed by Iterative Clustering with Guide Gene Selection, version 2 (ICGS2) to generate clusters and Uniform Manifold Approximation and Projection (UMAP) plots. We annotated the clusters based on surface marker phenotypes and gene expression (Fig. 2A, B and Supp. Fig. 3). Slingshot was used to infer differentiation trajectories, which appeared similar in wildtype and MLL::ENL-expressing cells irrespective of when MLL::ENL was induced (Fig. 2A). Clusters containing CD150^+^CD48^-^Lineage^-^Sca1^+^c-Kit^+^ HSCs and CD48^+^Lineage^-^Sca1^+^c-Kit^+^ MPPs (clusters 19 and 29, respectively) were underrepresented after fetal MLL::ENL induction, relative to the control group (Fig. 2C). Likewise, clusters containing Lineage^-^Sca1^-^c-Kit^+^CD16/32^-^ pre-Granulocyte-Monocyte Progenitors (pGMs) and Lineage^-^Sca1^-^c-Kit^+^CD16/32^+^ GMPs – clusters 12, 22 and 9 – were overrepresented after fetal MLL::ENL induction (Fig. 2C). These shifts were not observed after postnatal induction (Fig. 2D). MLL::ENL expression was lower after fetal induction than after postnatal induction, consistent with bulk RNA-seq results (Fig. 2E). *Hoxa9* expression was similar after fetal and postnatal MLL::ENL induction, demonstrating that even with lower transgene expression, some MLL::ENL targets remain overexpressed (Fig. 2F). Altogether, the data show that fetal MLL::ENL induction leads to a postnatal reduction in HSC/MPP frequency, increased myeloid differentiation and selection against cells with high oncogene expression.

**Figure 2.**
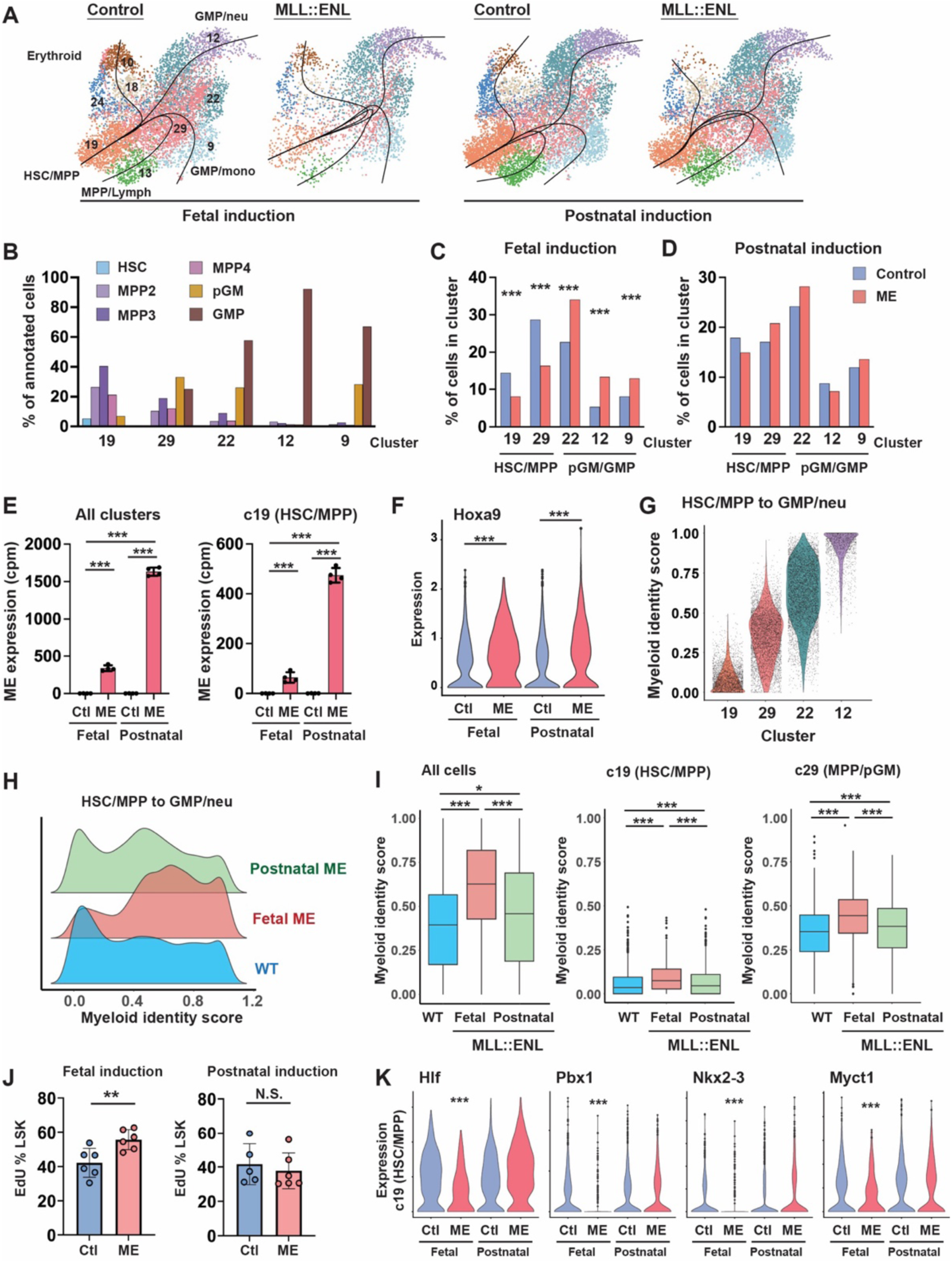
Fetal MLL::ENL induction enhances postnatal myeloid priming. (A) UMAPs representing single cell gene expression in P28 LK cells after fetal or postnatal MLL::ENL (ME) induction. Black lines reflect differentiation trajectories as calculated by Slingshot. (B) Cell identities by cluster based on surface marker phenotypes. (C, D) Distribution of cells within indicated clusters of P28 LK progenitors following fetal or postnatal MLL::ENL induction. ***p<0.001 by Wicoxon signed-rank test. (E) *MLL::ENL* transgene expression in all clusters (left panel) or the HSC/MPP cluster (c19, right panel) based on alignment of the human *MLL::ENL* transcript in n=4 pseudoreplicates. ***p<0.001 by Wilcoxon signed-rank test. (F) *Hoxa9* gene expression in control and MLL::ENL HSC/MPPs (cluster 19) ***p<0.001 by Wilcoxon signed-rank test. (G) Quadratic programming-derived myeloid identity scores for clusters along the HSC/MPP to GMP-neu trajectory in P28 wildtype mice. (H) Histogram showing distribution of myeloid identity score in all cells along the HSC/MPP to GMP-neu trajectory after fetal or postnatal MLL::ENL induction. (I) Distribution of myeloid identity score in all cells along the HSC/MPP to GMP-neu trajectory, HSC/MPPs, and MPP/pGMs after fetal or postnatal MLL::ENL induction. *p<0.05, ***p<0.001 by Wilcoxon signed-rank test. (J) EdU incorporation for P28 LSKs and GMPs following E10.5 MLL::ENL induction. **p<0.01 by student’s t-test. (K) Expression of self-renewal genes in HSC/MPPs after fetal or postnatal MLL::ENL induction. ***p<0.001 relative to age matched control by Wilcoxon signed-rank test.

The relative reduction in HSC/MPPs after fetal MLL::ENL induction raised the question of whether this reflects enhanced myeloid priming within one or more progenitor populations. To test this possibility, we used quadratic programming to assign myeloid identity scores to each cluster along the HSC to granulocyte-monocyte progenitor/neutrophil (GMP/neu) differentiation trajectory (Fig. 2A, G). Each cell was assigned a score between 0 and 1 based on expression of genes that distinguished the HSC/MPP cluster 19 from the GMP/neu cluster 12 in wildtype P28 mice (Fig. 2G) (25). Fetal MLL::ENL induction resulted in higher myeloid identity scores, relative to the controls, across the entire differentiation trajectory, including HSC/MPP and MPP/pGM clusters (Fig. 2H, I). Postnatal MLL::ENL induction also drove a statistically significant increase in myeloid differentiation but to a far less extent than was observed after fetal induction (Fig. 2I). Thus, fetal MLL::ENL induction enhances myeloid priming after birth, even in HSCs and MPPs.

We next tested whether fetal and postnatal MLL::ENL induction have distinct effects on global gene expression and cell proliferation at P28. We performed pseudobulk gene expression analysis for HSC/MPP and pGM/GMP clusters, followed by Gene Set Enrichment Analysis (GSEA) and Gene Set Variance Analysis (GSVA) to identify gene signatures that that distinguish fetal and postnatal induction cohorts (Supp. Table 1). MYC targets, E2F targets, mTORC1-associated genes and oxidative phosphorylation-associated genes were all elevated after fetal but not postnatal MLL::ENL induction, particularly in the HSC/MPP cluster (Supp. Fig. 4A-C). In addition, a higher percentage of HSC/MPP cells appeared to be in S-phase after fetal MLL::ENL induction than after postnatal induction (Supp. Fig. 4D). EdU (5-ethynyl-2’-deoxyuridine) incorporation assays confirmed that P28 HSC/MPPs cycle more frequently after fetal MLL::ENL induction but not postnatal MLL::ENL induction (Fig. 2J). MLL::ENL did not alter GMP proliferation at either age (Supp. Fig. 4E). Fetal MLL::ENL induction significantly reduced expression of several HSC self-renewal genes within the HSC/MPP cluster, including *Hlf*, *Pbx1*, *Nkx2-3* and *Myct1* (Fig. 2K, Supp. Fig. 4F). This was not observed after postnatal induction. Thus, fetal MLL::ENL induction leads to postnatal transcriptional changes in HSC/MPPs that reflect enhanced myeloid priming, reduced self-renewal and elevated rates of proliferation. These changes were evident despite selection against cells that most highly expressed MLL::ENL.

### MLL::ENL drives precocious myeloid gene expression without corresponding changes in chromatin architecture

We next tested whether MLL::ENL elicits distinct changes in chromatin organization after fetal and postnatal induction, given that fetal resistance to AML initiation is heritable and potentially sustained through epigenome remodeling. We induced MLL::ENL before or after birth and isolated LK cells at P28 for single-cell Assay for Transposase-Accessible Chromatin using sequencing (scATAC-seq). The cells were clustered using ArchR. CITE-seq data were then integrated to annotate clusters based on gene expression and to assess concordance between differentiation states, as defined by chromatin accessibility and gene expression (Fig. 3A, B). scATAC-seq cluster distributions were surprisingly similar after fetal and postnatal MLL::ENL induction (Fig. 3A, C). However, there was clear discordance between the scATAC-seq and CITE-seq cluster assignments. For example, scATAC-seq clusters 6 and 3 each had cells assigned to CITE-seq clusters 22 (pGM) or 12 (GMP-neu) based on integrated transcript expression. A lower percentage of cells in each scATAC-seq cluster were defined, transcriptionally, as pGM after fetal MLL::ENL induction than after postnatal induction, and a greater percentage were defined as GMP-neu (Fig. 3D). We used quadratic programming to quantify this discrepancy (Fig. 3E). For every cluster of cells with similar degrees of epigenetic differentiation, the degree of myeloid bias, as defined by integrated transcript expression, was significantly higher after fetal MLL::ENL induction than after postnatal induction (Fig. 3E). Thus, MLL::ENL promotes precocious myeloid differentiation after fetal induction without altering chromatin accessibility.

**Figure 3.**
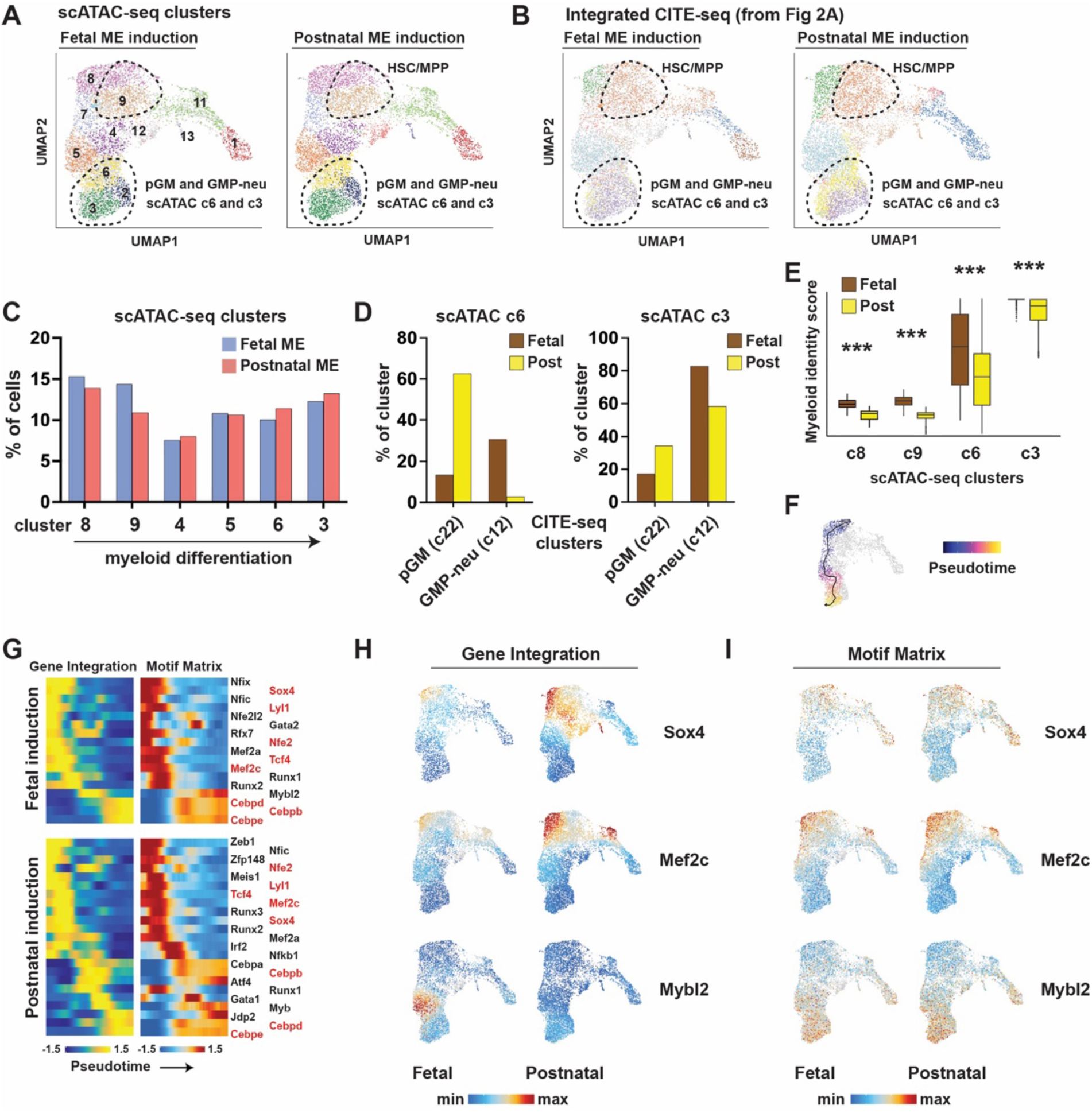
Fetal MLL::ENL induction provokes myeloid-biased gene expression without altering chromatin accessibility. (A) UMAP representing control and MLL::ENL-expressing (ME) P28 LK cells based on scATAC-seq profiles. Clustering was performed in ArchR. (B) Cluster assignments based on inferred gene expression. The assignments follow the clustering nomenclature specified in Figure 2A. (C) Distribution of cells from fetal and postnatal MLL::ENL induction groups within scATAC-seq clusters. (D) Inferred CITE-seq identities for cells within scATAC-seq clusters 3 and 6. (E) Myeloid identity scores, based on quadratic programming using integrated CITE-seq data, for cells within the indicated scATAC-seq clusters after fetal or postnatal MLL::ENL induction. ***p<0.0001 by Wilcoxon signed rank test. (F) Pseudotime trajectory used to define coordinated changes in transcription factor expression and motif accessibility during myeloid differentiation. (G) Transcription factors with concordant differential expression (based on integrated CITE-seq data) and motif accessibility (based on scATAC-seq) through the course of myeloid differentiation, after fetal or postnatal MLL::ENL induction. Genes in red were identified in both cohorts. (H) *Sox4*, *Mef2c* and *Mybl2* transcript expression in individual cells, projected as heatmaps on the UMAPs from panel A. Ranges of min/max expression are identical for fetal and postnatal induction cohorts for each gene. (I) Enrichment for SOX4, MEF2C and MYBL2 motifs within individual cells, projected as heatmaps.

We next compared differentiation programs of MLL::ENL-expressing progenitors after fetal and postnatal induction. We plotted a myeloid differentiation trajectory in pseudotime based on scATAC-seq clusters (Fig 3F). Using ArchR and ChromVar, we identified transcription factors that 1) were differentially expressed through the course of myeloid differentiation based on integrated CITE-seq data, and 2) had corresponding changes in binding site accessibility through the course of differentiation (Fig. 3G). There was partial overlap between transcription factors that met these criteria after fetal and postnatal MLL::ENL induction (Fig. 3G), but inspection of UMAP plots showed highly variable transcription factor expression between the two induction ages (Fig. 3H, Supp. Fig. 5A). For example, two transcription factors that have known roles in maintaining stemness, *Sox4* and *Mef2c* (26,27), were expressed in both induction groups, but more diffusely and at higher levels after postnatal induction (Fig. 3H). In contrast, *Mybl2*, a transcription factor with an established role in myeloid differentiation (28), was more highly and diffusely expressed after fetal MLL::ENL induction (Fig. 3H). Several other transcription factors showed distinct patterns of expression after fetal and postnatal induction (Supp. Fig. 5A). For all of these transcription factors, binding site accessibility patterns were largely identical between the two groups (Fig. 3I, Supp. Fig. 5B). These data again underscore the profound discordance between effects of MLL::ENL on gene expression and chromatin accessibility. They imply that the heritable fetal barrier to AML is enforced at the level of transcriptional regulation rather than through epigenome reorganization.

### MLL::ENL suppresses HSC/MPP proliferation during late-gestation and alters gene expression without chromatin remodeling

We next evaluated MLL::ENL-dependent changes in gene expression and cell proliferation during the late fetal and early neonatal stages of development, given that the selective pressure against MLL::ENL expression is established during this time period. We first performed CITE-seq on P0 LK progenitors isolated from livers of Tet-Off-ME mice after fetal MLL::ENL induction. We clustered and annotated cells based on gene expression and surface marker phenotypes (Fig. 4A; Supp. Fig. 6A). In contrast to what was observed at P28, fetal MLL::ENL induction did not lead to precocious myeloid differentiation at P0. HSC, MPP and GMP cluster frequencies, and myeloid identity scores, were similar between the control and MLL::ENL-expressing samples even with robust MLL::ENL induction (Fig. 4B-D). GSEA showed reduced expression of genes associated with cell proliferation within the MLL::ENL-expressing MPP cluster along with increased expression of genes associated with type-I Interferon (IFN-1) signaling (Fig. 4E, F; Supp. Table 2). EdU incorporation assays, performed at E18, confirmed that MLL::ENL suppresses HSC/MPP proliferation during late gestation, though GMP proliferation was unaffected (Fig. 4G, Supp. Fig. 6B). Deleting the interferon alpha receptor 1 (*Ifnar1*) had no effect on AML initiation after fetal MLL::ENL induction, despite enrichment for the IFN-1 signature (Fig. 4H). Thus, MLL::ENL suppresses HSC/MPP proliferation during late gestation, but perinatal interferon signaling does not impede leukemogenesis. Reduced HSC/MPP proliferation could convey a selective disadvantage that ultimately favors cells with lower levels of MLL::ENL expression.

**Figure 4.**
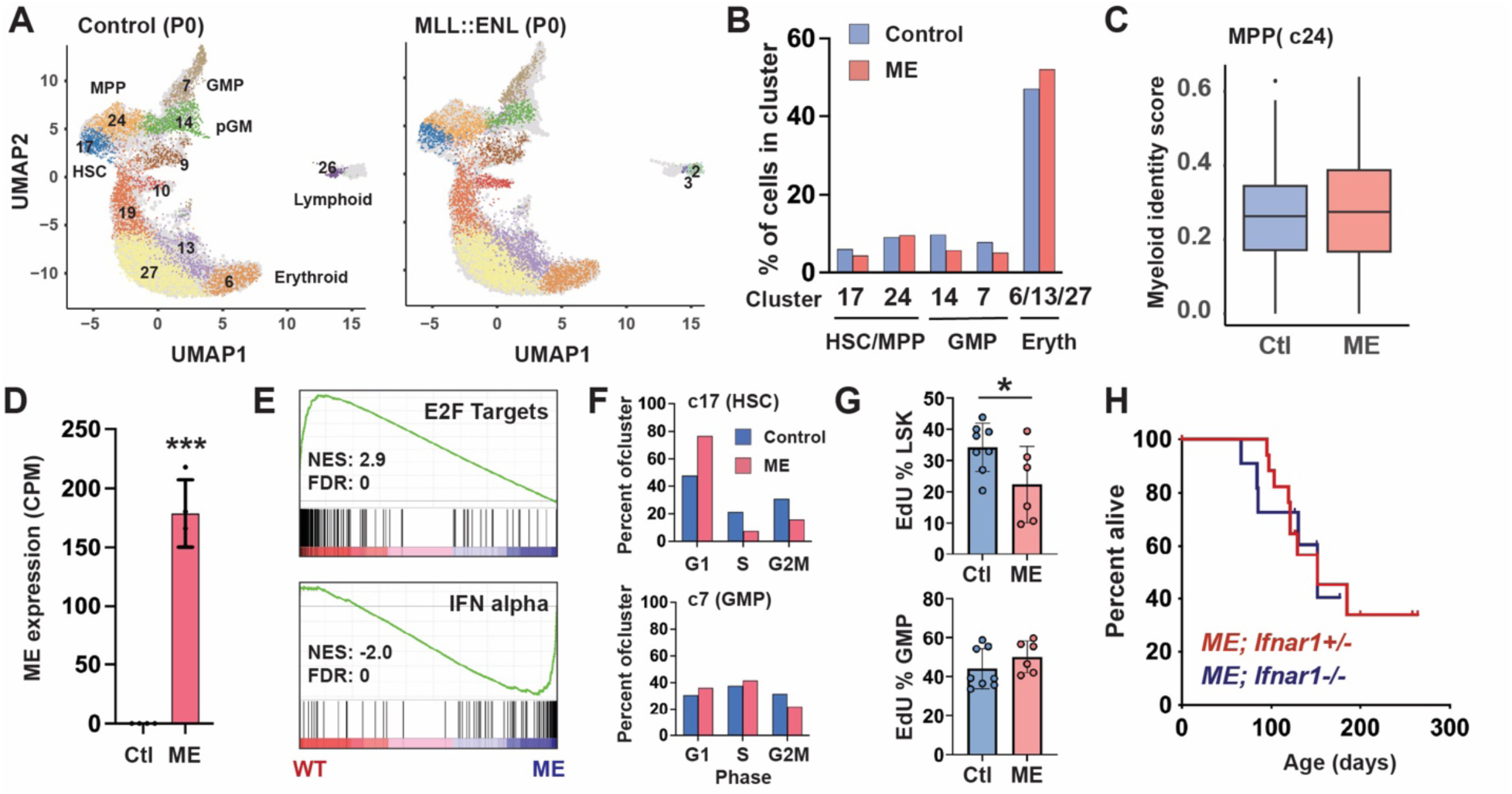
Fetal MLL::ENL induction suppresses HSC/MPP proliferation during late-gestation. (A) UMAP representing P0 LK cells, cluster identities and annotation based on CITE-seq performed after fetal MLL::ENL induction. (B) Cluster distributions from control and MLL::ENL-expressing (ME) cohorts. (C) Myeloid identity scores for cells in MPP cluster 24, as calculated by quadratic programming. (D) *MLL::ENL* transgene expression in HSC cluster 17 based on n=4 pseudoreplicates. ***p<0.001. (E) GSEA plots show suppression of E2F target genes and activation of IFN alpha target genes after MLL::ENL induction, as calculated based on four pseudoreplicates for HSC cluster 17. (F) Percent of cells within HSC (c17) and GMP (c7) clusters at indicated cell cycle phases based on single cell gene expression. (G) EdU incorporation in E18 LSK and GMP cells. n= 8 mice for control and n= 6 mice for Tet-Off-MLL::ENL samples, *p<0.05 by two-tailed Student’s t-test. (H) Kaplan-Meier survival curves for Tet-ON-ME; *Ifnar1*^+/-^ and Tet-ON-ME; *Ifnar1*^-/-^ mice after E10.5 MLL::ENL induction.

To assess mid rather than late gestational changes in gene expression, we transitioned to a novel Tet-On approach that enabled us to isolate and analyze hematopoietic progenitors more efficiently than conventional breeding strategies. We generated induced pluripotent stem cells (iPSCs) with *Vav1-Cre*, *Col1a1^FRT^* and *Rosa26^CAG-LSL-rTTA-IRES-mKate2^*alleles (Fig. 5A). The *Col1a1^FRT^* allele provides a safe harbor locus for FLP-mediated insertion of DOX-regulated transgenes. We inserted a 3x-FLAG-MLL::ENL transgene into this locus. The remaining alleles were identical to those used in the Tet-On-ME model shown in Figure 1. We used the iPSCs to generate chimeric mice, such that blood ontogeny proceeded normally (Fig. 5A). iPSC-derived hematopoietic cells could be identified based on mKate2 expression. This strategy served two purposes. First, a Tet-On system ensured precise induction of MLL::ENL upon DOX exposure during a tight gestational window, whereas Tet-Off systems would have required gradual DOX clearance. Second, all chimeric mice generated with this system had the same Tet-On-ME genotype in iPSC-derived cells. This greatly simplified efforts to collect cells from embryos with complex genetic backgrounds for molecular studies. As with the previously described models, iPSC-derived Tet-On-ME progenitors exhibited robust MLL::ENL expression at E18 after induction at E10.5, but protein expression declined after birth (Fig. 5B). Furthermore, when we transplanted chimeric LK cells after fetal or postnatal MLL::ENL induction, AML onset in recipients was delayed and less penetrant after fetal induction as compared to postnatal induction (Fig. 5C). Chimeric mice therefore exhibit the same fetal barrier to MLL::ENL-driven leukemogenesis as was observed in non-chimeric Tet-On-ME and Tet-Off-ME mice.

**Figure 5.**
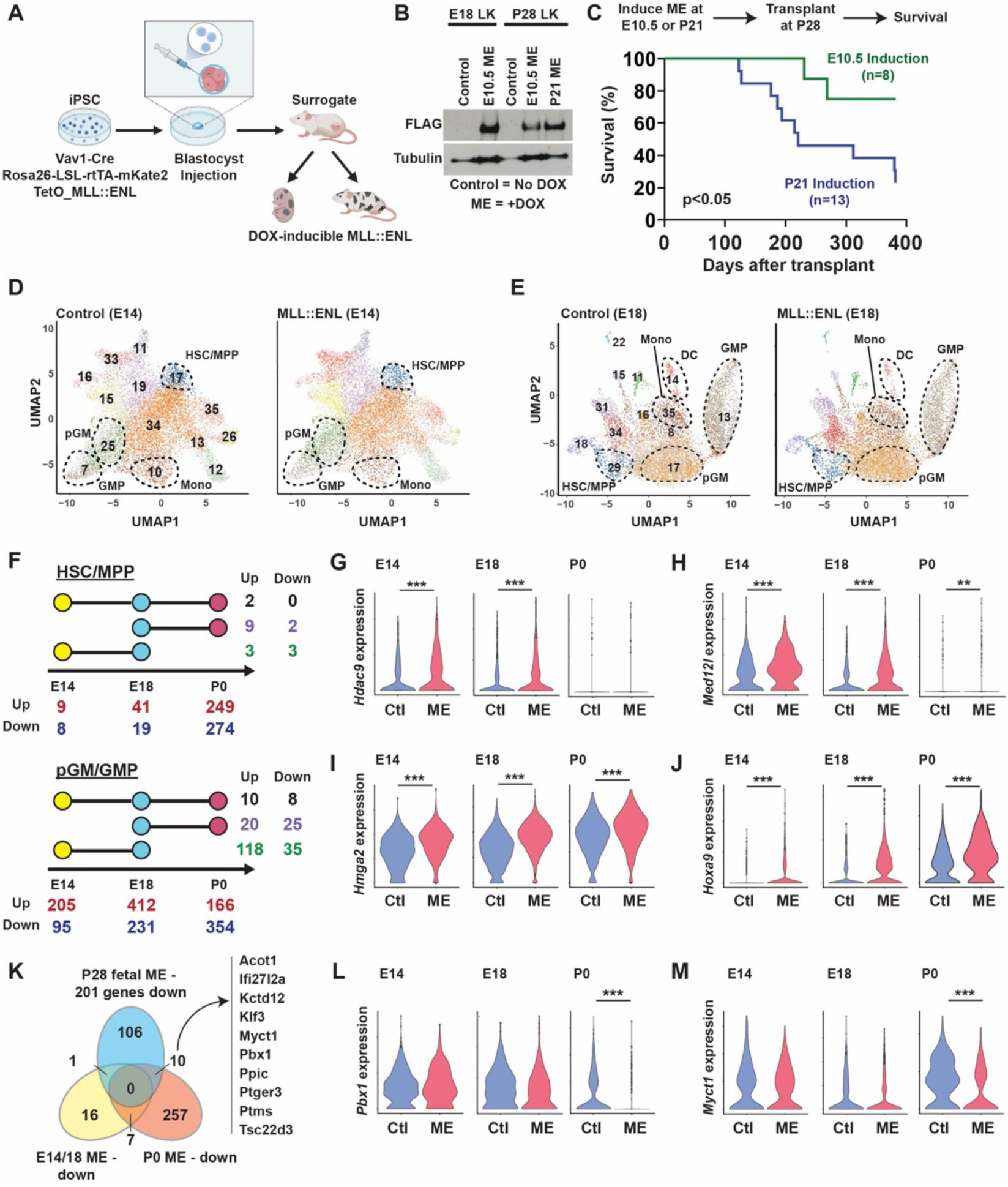
Fetal MLL::ENL induction drives changes in gene expression at mid-gestation that are not sustained during late-gestation and postnatal stages. (A) Overview of iPSC-derived chimeric, DOX-inducible MLL::ENL mouse model. (B) Western blot showing expression of FLAG-tagged MLL::ENL in E18 LSK following fetal induction, and P28 LSK following fetal or postnatal induction. (C) Kaplan-Meier survival curve for recipients of iPSC-derived, MLL::ENL-expressing LSK cells following induction at E10.5 or P21. (D, E) UMAPs representing CITE-seq profiles for iPSC-derived, E14 or E18 LK cells after MLL::ENL induction at E10.5. Annotation is based on gene expression and surface marker expression (Supp. Fig. 7A, B). Control cells were derived from chimeric mice that were not administered DOX. (F) Summary of the number of significantly upregulated or downregulated genes in HSC/MPP (top panel) and pGM/GMP (bottom panel) cells from CITE-seq data performed at E14, E18, and P0. Significant differential expression was defined as log2(fold change) >0.6 and p.adj. <0.05 based on 4 pseudoreplicates for each cluster. Total numbers of upregulated and downregulated genes for each age/cell type are shown beneath the plots. Connected circles indicate overlap among profiles of differentially expressed genes across the ages, with numbers shown to the right of the plots. (G-J) Expression of *Hdac9* (G), *Med12l* (H), *Hmga2* (I), and *Hoxa9* (J) at indicated ages. **p<0.01, ***p<0.0001 by Wilcoxon signed-rank test. (K) Venn diagram summarizing genes that are significantly downregulated – log2(fold change) >0.6 and P.adj. <0.05 – in MLL::ENL expressing HSC/MPP clusters, relative to age-matched controls, at fetal (E14 and E18), neonatal (P0), and postnatal (P28) stages. (L, M) Expression of *Pbx1* (L) and *Myct1* (M) at indicated ages. ***p<0.0001 by Wilcoxon signed-rank test.

We performed CITE-seq on E14 and E18 chimeric LK cells after inducing MLL::ENL at E10.5. We identified clusters containing HSC/MPP cells, pGMs and GMPs based on surface marker phenotypes (Fig. 5D, E; Supp. Fig. 7A, B). MLL::ENL expression did not dramatically alter distributions of cells within these clusters at either age (Supp. Fig. 7C, D). MLL::ENL had only limited immediate effects on HSC/MPP gene expression at E14, though the number of differentially expressed genes increased with gestational age (Fig. 5F). Changes that did occur at E14, such as *Hdac9* and *Med12l* induction, were not sustained at P0 (Fig. 5G, H). MLL::ENL had far greater effects on gene expression in E14 pGM/GMP clusters, though again, only a small number of changes were sustained at P0 (Fig. 5F). Only 10 of the 166 significantly upregulated genes, and 20 of the 354 significantly downregulated genes observed at P0 were also differentially expressed at E14 or E18. These included know targets of MLL fusion proteins - *Hmga2*, *Hoxa5*, *Hoxa7* and *Hoxa9* (Fig. 5I, J, Supp. Table 3, 4). Only a small number of genes that were downregulated in P28 HSC/MPPs after fetal induction (relative to postnatal induction) were already downregulated at P0 (Fig. 5K). These included the HSC self-renewal genes *Myct1* and *Pbx1*, neither of which were significantly downregulated before birth (Fig. 5L, M). These comparisons lead to the overarching conclusion that many of the changes in gene expression that become evident at or after birth following fetal MLL::ENL induction are secondary or tertiary consequences of fetal oncoprotein exposure. Very few of the changes that occur immediately after fetal MLL::ENL induction are actually sustained. This supports a scenario of selection against MLL::ENL expressing cells through the course of late fetal and early neonatal ontogeny.

We tested whether MLL::ENL drives changes in chromatin accessibility before birth or whether there is discordance between transcriptional and epigenomic cluster assignments, as was observed at P28. We performed scATAC-seq on E18 mKate+ LK cells harvested from chimeric mice after inducing MLL::ENL at E10.5. We used ArchR to assign cluster identities based on chromatin accessibility and integrated E18 CITE-seq data (Fig. 6A, B). MLL::ENL expression did not cause marked redistribution of cells within the scATAC-seq clusters (Fig. 6C). Moreover, we did not observe discordance between scATAC-seq and CITE-seq cluster assignments at E18 (Fig. 6D). For example, almost all cells in scATAC-seq clusters 13 and 15 were assigned to CITE-seq cluster 29 (HSC/MPP), and a majority of cells in scATAC-seq cluster 9 were assigned to CITE-seq cluster 13 (GMP) (Fig. 6D). MLL::ENL expression had minimal effect on these distributions. Thus, fetal MLL::ENL induction does not lead to discordant myeloid transcriptional priming until after birth.

**Figure 6.**
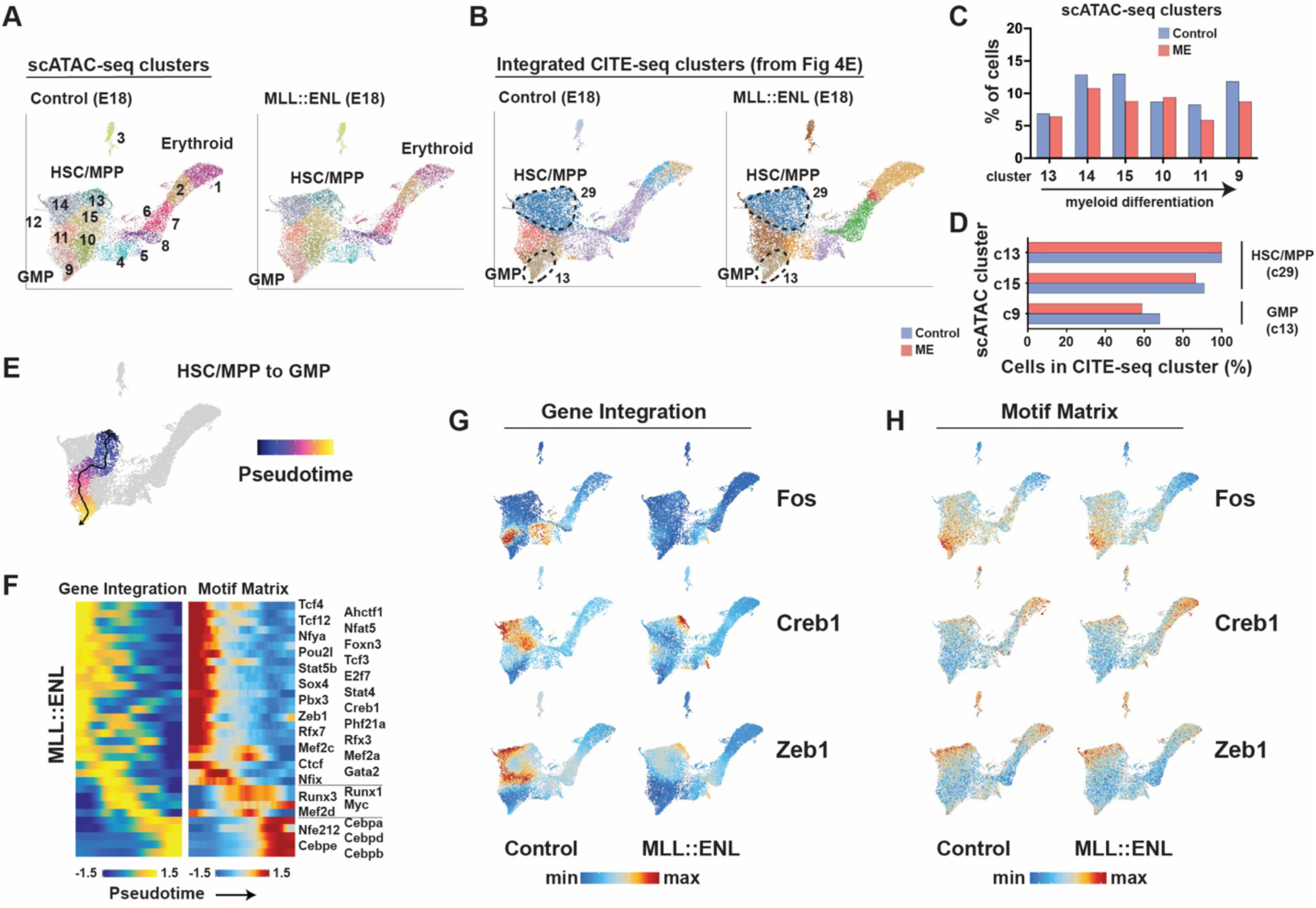
MLL::ENL alters transcription factor expression but not chromatin organization during fetal hematopoiesis. (A) UMAPs representing scATAC-seq profiles for iPSC-derived E18 LK cells after MLL::ENL induction at E10.5. (B) Cluster assignments based on inferred gene expression. The assignments follow the clustering nomenclature specified in Figure 5E. (C) Distribution of cells within scATAC-seq clusters along the myeloid differentiation trajectory. (D) Percentages of cells with HSC/MPP and GMP cluster assignments, based on inferred transcript expression, in scATAC-seq clusters 9, 13 and 15. (E) Pseudotime trajectory used to define coordinated changes in transcription factor expression and motif accessibility during myeloid differentiation. (F) Transcription factors with concordant differential expression (based on integrated CITE-seq data) and motif accessibility (based on scATAC-seq) through the course of myeloid differentiation in MLL::ENL expressing E18 LK cells. (G) *Fos*, *Creb1* and *Zeb1* transcript expression in individual cells, projected as heatmaps on the UMAPs from panel A. Ranges of min/max expression are identical for fetal and postnatal induction cohorts for each gene. (H) Enrichment for FOS, CREB1 and ZEB1 motifs within individual cells, projected as heatmaps.

We next identified transcriptions factors that exhibit changes in expression and motif accessibility through the course of differentiation by plotting a differentiation trajectory in pseudotime (Fig. 6E, F). Many transcription factors exhibited discrepant expression between control and MLL::ENL-expressing progenitors (Fig. 6G; Supp. Fig. 8A). These included *Fos*, *Junb*, *Runx1*, *Mef2a*, *Creb1* (a critical driver of leukemic transformation)(29) and *Zeb1* (a critical HSC self-renewal gene)(30) (Fig. 6G; Supp. Fig. 8A). All of these genes were expressed at either lower levels or across a smaller population of cells after MLL::ENL induction, yet changes in motif accessibility were not observed (Fig. 6H; Supp. Fig. 8B). These data show that MLL::ENL alters the complement of transcription factors that are expressed during fetal hematopoietic differentiation without causing chromatin reorganization, ultimately yielding an AML resistant transcriptional state.

### MLL3 enforces AML resistance in myeloid primed, MLL::ENL-expressing progenitors

We next sought mechanisms that maintain the fetal barrier to AML initiation after birth and transplantation. The lack of MLL::ENL-dependent changes in chromatin accessibility argued against chromatin remodeling factors, but the heritable nature of the resistance argued for an epigenetic mechanism. We focused our attention on MLL3, a histone methyltransferase that promotes HSC and GMP differentiation but does not modulate bulk ATAC-seq profiles of HSCs, MPPs or GMPs (31). We previously showed that MLL3 suppresses *Hox* gene expression in GMPs, suggesting that it may have a role in suppressing *MLL*r AML (32). Furthermore, *MLL3* is somatically mutated in pediatric AML (10), and *MLL3* mutations have been shown to convey resistance to MENIN inhibitors, which are used to treat *MLL*r leukemias (33). Prior studies have shown that MLL3 binds cis-regulatory elements and promotes gene expression by facilitating RNA polymerase II pause-release (34). Together, these observations raised the question of whether MLL3 helps maintain an AML resistant state after fetal MLL::ENL induction.

To assess whether *Mll3* maintains the fetal AML barrier, we crossed Tet-Off-ME mice to previously described *Mll3^f/f^* mice and tested whether *Mll3* deletion could enable transformation after fetal MLL::ENL induction (31,32). We induced MLL::ENL at E10.5 and transplanted P0 liver progenitors as in Fig. 1E. Almost all recipients of *Mll3^+/+^* LK cells survived greater than 200 days post-transplant (Fig. 7A). In contrast, all recipients of *Mll3*-deficient (*Mll3*^Δ/Δ^) cells died with a median survival of 90 days (Fig. 7A), indicating a role for MLL3 in sustained AML resistance. We next crossed Tet-Off-ME mice to *Mll3^+/-^*mice and evaluated survival after fetal or postnatal MLL::ENL induction without performing transplants. As in prior experiments, fetal MLL::ENL induction on an *Mll3^+/+^* background led to AML but with incomplete penetrance and long latency (Fig. 7B). Deleting just a single *Mll3* allele greatly accelerated AML onset (Fig. 7B). *Mll3* deletion had only a modest effect on leukemogenesis after postnatal MLL::ENL induction, as might be expected given the rapid AML onset that was observed even on an *Mll3^+/+^* background (Fig. 7C). Altogether, these data show that MLL3 is necessary to maintain resistance to AML initiation that ensues after fetal MLL::ENL initiation.

**Figure 7.**
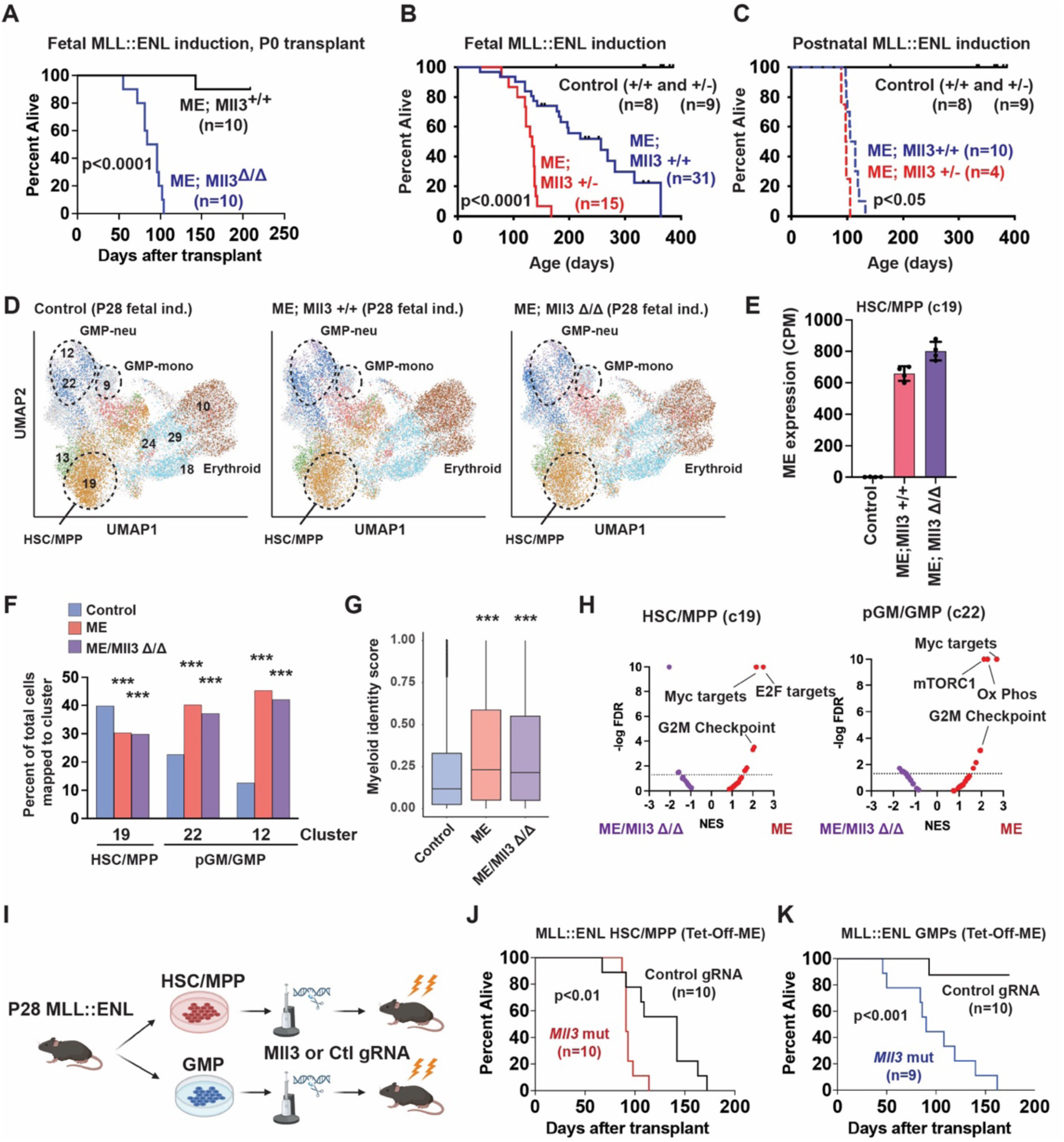
*Mll3* sustains a leukemia resistant state in myeloid primed MLL::ENL-expressing postnatal progenitors. (A) Kaplan-Meier survival curves for recipients of Tet-Off-ME; *Mll3*^Δ/Δ^ P0 liver cells after fetal MLL::ENL induction. (B, C) Kaplan-Meier survival curves for Tet-Off-ME; *Mll3^+/+^* and Tet-Off-ME; *Mll3^+/-^* mice after fetal or postnatal MLL::ENL induction. (D) UMAPs representing CITE-seq profiles for P28 LK cells from control, Tet-Off-ME; *Mll3^+/+^* and Tet-Off-ME; *Mll3*^Δ/Δ^ mice. Cluster annotations were assigned based on Symphony alignment to the data shown in Figure 2A. (E) *MLL::ENL* transgene expression in HSC/MPP cluster 19. (F) Percentages of cells for each genotype in clusters 19, 22 and 12. (G) Distribution of myeloid identity scores in all cells along the HSC/MPP to GMP-neu trajectory, as calculated by quadratic programming. ***p<0.001 by Wilcoxon signed-rank test. (H) Volcano plots showing normalized enrichment scores and false discovery rates for Hallmark gene sets, as calculated from 4 pseudoreplicates per genotype for the HSC/MPP and pGM/GMP populations. (I) Approach to mutagenize *Mll3* in HSC/MPPs and GMPs from P28 Tet-Off-ME mice. (J, K) Kaplan-Meier survival curves for recipients of Tet-Off-ME HSC/MPPs (J) or GMPs (K) after targeted mutagenesis of an intragenic region on chromosome 8 or *Mll3*. (K) For all survival curves, group sizes and p-values are shown in the panels. P-values are indicated and were calculated by the log rank test.

To determine how MLL3 contributes to sustained AML resistance, we performed CITE-seq on control (*Vav1-Cre* negative), Tet-Off-ME; *Mll3^+/+^* and Tet-Off-ME; *Mll3^Δ/Δ^* LK progenitors harvested at P28 after MLL::ENL induction at E10.5. UMAPs were generated with ICGS2, as in the prior experiments, but in this case, we used Symphony to assign cells to clusters using the same identities as were assigned in Figure 2A (Fig. 7D). This approach allowed us to evaluate changes in cluster distribution and myeloid priming consistently across the different CITE-seq experiments. We evaluated expression of the MLL::ENL transcript in HSC/MPP cluster 19. Expression was comparable between the *Mll3^+/+^* and *Mll3^Δ/Δ^* populations (Fig. 7E). Thus, *Mll3* deletion does not relieve the selective pressure that leads to lower postnatal MLL::ENL expression, even though it greatly accelerates leukemogenesis. MLL::ENL-dependent myeloid priming was evident in both *Mll3^+/+^*and *Mll3^Δ/Δ^* progenitors, relative to wildtype progenitors (Fig. 7F, G; Supp. Table 5). *Mll3* deletion did reverse enrichments in MYC, mTORC1 and E2F target gene sets that were observed after MLL::ENL induction (Fig. 7H). The overarching conclusion drawn from these data is that MLL3 restricts the capacity of myeloid primed cells to transform after fetal MLL::ENL induction, rather than enforcing the myeloid bias itself.

In light of these findings, we tested whether *Mll3* deficiency enables efficient transformation of MLL::ENL-expressing GMPs. Prior work has shown that MPPs, pGMs and GMPs can all serve as a cell of origin for MLL::ENL-driven AML, but GMPs are much less efficiently transformed (35). We induced MLL::ENL postnatally in Tet-Off-ME mice and isolated HSC/MPPs (LSK cells) or GMPs at P28. We used CRISPR/Cas9 to mutate *Mll3* and transplanted 4000 cells per recipient into lethally irradiated recipient mice (Fig. 7I). As expected for the postnatal induction age, all recipients of Tet-Off-ME HSC/MPPs developed AML, though *Mll3* deletion significantly reduced survival time (Fig. 7J). Very few recipients of control Tet-Off-ME GMPs developed AML, whereas all recipients of *Mll3* mutated GMPs succumbed to AML (Fig. 7K). Thus, MLL3 enforces an AML resistant state in myeloid committed progenitors.

## DISCUSSION

Our data offer a potential explanation for why congenital leukemias are exceedingly rare despite high rates of prenatal progenitor expansion, documented prenatal acquisition of *MLL*r and low mutation thresholds required for transformation. MLL::ENL imposes a negative selective pressure that is much stronger when the mutation arises before birth rather than after birth. The fusion protein suppresses fetal HSC/MPP proliferation to an extent not seen after birth (Fig. 3). This ultimately leads to postnatal outgrowth of progenitors that maintain only low levels of MLL::ENL expression and that lose critical stemness programs in favor of precocious myeloid differentiation (Fig. 2). While corresponding events are difficult to resolve in human fetuses, the murine data suggest that when an *MLL*r occurs during human fetal hematopoiesis, affected progenitors may be outcompeted by normal cells and differentiate towards stable, non-malignant fates. This model runs counter to most prevailing models of tumorigenesis, wherein sequentially acquired mutations convey selective advantages rather than selective disadvantages (36). Furthermore, it adds a layer of nuance to prior models that argued for enhanced leukemia susceptibility during fetal stages of life (18–20).

The data raise the question as to why *MLL*r congenital and infant leukemias occur at all, given that expression of an MLL fusion protein in fetal progenitors conveys a sustained, selective disadvantage. It is important to note that the fetal barrier to transformation is relative rather than absolute, and we have identified two mechanisms that can circumvent it. First, a recurrent cooperating mutation, *Nras*^G12D^, did enable highly penetrant transformation even after fetal MLL::ENL induction (Fig.1B, Supp. Fig. 1C). Second, *Mll3* deletion mitigated the barrier by extending efficient transformation capacity to more differentiated progenitors, thus expanding the repertoire of potential cells of origin (Fig. 7). Altogether, the data suggest that following a *MLL*r, fetal cells may have a narrow window of time to acquire cooperating mutations before they are outcompeted and lose leukemogenic potential. This makes infant and congenital leukemias rare, but not impossible.

It is still not clear why MLL::ENL elicits distinct transcriptional changes in fetal and postnatal progenitors. Many of the transcriptional changes that arose in newborn and juvenile progenitors after fetal MLL::ENL induction appear to be indirect changes, given that they were not evident during mid gestation. We were unable to obtain sufficient number of fetal progenitors to identify direct MLL::ENL interactions with chromatin. It therefore remains an open question as to whether MLL::ENL binds different target genes at different ages or whether the targets activate distinct secondary targets. Regardless, the mechanisms that account for age-specific responses to MLL::ENL are reinforced at the transcriptional level rather than through chromatin remodeling. This suggests that normal heterochronic transcription factors may interact with MLL::ENL targets, or MLL::ENL itself, to exert a negative selective pressure in fetal progenitors.

One of the most fascinating aspects of developmental biology is that single blastocysts can expand exponentially to become trillion celled organisms with near-perfect fidelity. The fidelity is not simply spatial and functional. It also reflects the absence of neoplasia in the setting of rapidly expanding, highly stimulated developmental fields. Our data suggest that low rates of in utero cancers may reflect active diversion of mutant cells towards non-malignant fates, in addition to the lower mutation burdens inherent to early life. Going forward, it will be important to establish whether such barriers apply to other cancer-causing mutations and why they dissipate shortly after birth.

## METHODS

### Mouse lines

*Vav1-Cre* (RRID:IMSR_JAX:008610), *Rosa26^LSL-rtTA-IRES-mKate2^*(RRID:IMSR_JAX:008610), *Rosa26^LSL-tTA^* (RRID:IMSR_JAX:011008), *Col1a1^TetO-H2B-GFP^* (RRID:IMSR_JAX:016836) and *Nras^G12D^* (RRID:IMSR_JAX:008304), were obtained from Jackson Laboratory. *Col1a1^TetO-MLL::ENL^* (from David Bryder, Lund) and *Kmt2c^flox^* mice (RRID:IMSR_JAX:037115) were described previously (31,35). DOX chow (200 ppm) was purchased from Bioserv. Transplantations were performed by retro-orbital injections.

iPSC-derived chimeric mice were generated by first interbreeding mice with *Vav1-Cre*, *Col1a1^FRT^* and *Rosa26^LoxP-STOP-LoxP-rTTA-mKate2^* alleles. The *Col1a1^FRT^* was in derived from *Col1a1^TetO_H2B-GFP^* by crossing with Bact-Flpe mice (JAX) to remove the H2B-GFP cassette and restore a single functional FRT site in the *Col1a1* locus. Compound *Vav1-Cre*; *Col1a1^FRT^; Rosa26^LoxP-STOP-LoxP-rTTA-mKate2^* mouse embryonic fibroblasts were isolated and reprogrammed into iPSCs using the CytoTune-iPS 2.0 Sendai Reprogramming Kit (Invitrogen, A16517). iPSCs were nucleofected with pCol-MLL::ENL and AAV-Flpe constructs and placed under hygromycin selection. Successfully targeted clones were confirmed by PCR. Cells were then karyotyped and stained for SOX2 and OCT4 expression to confirm pluripotency. iPSCs were injected into blastocysts and implanted into pseudopregnant CD1 mice. iPSC-derived cells were isolated from fetal liver or neonatal bone marrow based on mKate2 expression.

### Study approval

All procedures were performed according to an Institutional Animal Care and Use Committee (IACUC)-approved protocol at Washington University School of Medicine.

### Sex as a biological variable

Both male and female mice were used in this study and analyzed together within the same groups.

### Flow cytometry

Cells were isolated, stained, analyzed, and transplanted as previously described (20,37,38). All antibodies were from Biolegend.

### RNA-seq library construction, sequencing, and analysis

Ten thousand LSK cells were isolated from P0 liver, neonatal bone marrow or transplant recipient bone marrow by flow cytometry. RNA was isolated with RNAeasy micro-plus columns (QIAGEN). Libraries were generated with Clontech SMRTer kits and sequenced on a Novaseq X plus. Sequences were aligned to the mouse genome using STAR version 2.0.4b. The sequence for human MLL::ENL was included in the alignment so that transgene transcript levels could be identified and quantified as counts per million within the pseudoreplicates.

### CITE-seq library construction, sequencing, and analysis

Bone marrow or fetal/neonatal liver cells were stained with APC-conjugated anti-CD117 and biotin-conjugated anti-Sca1 followed by anti-APC magnetic bead selection over Miltenyi columns. Two million Kit-selected cells were then stained with tagged Total-Seq B antibodies (Biolegend) and fluorescently tagged antibodies for flow cytometry. Sorted cells were loaded into Chromium Next GEM Chip G lanes (10x Genomics), and libraries were generated using Chromium Next GEM Single Cell 3′ kit v3.1 (PN-1000268), Chromium Single Cell 3’ Feature Barcode Library kit (PN-1000079), Dual Index kit NT Set A (PN-1000242) and Dual Index kit TT Set A (PN-1000215) reagents (all 10x Genomics). cDNA libraries were quantified on an Agilent Bioanalyzer and sequenced on an Illumina Novaseq X plus.

### CITE-seq analysis

The Cell Ranger v6.1.2 pipeline was used to align, filter and normalize digital gene expression files as previously described (37,38). We performed Iterative Clustering and Guide-gene Selection (ICGS, version 2) analysis with the AltAnalyze toolkit (39) to visualize cells and annotate clusters as previously described (37,38). For Figure 7, we used Symphony to assign cells to clusters using the same cluster identities from CITE-seq experiments performed in Figure 2 (40). For differential expression analyses, Presto was used to generate pseudoreplicates within clusters (4 per sample group) by aggregating read counts within each replicate (41). DESeq2 was then used to identify differentially expressed genes (42). Pseudoreplicates were also used to perform to perform GSEA (43). GSVA was performed using a Gaussian kernel as previously described (37,44). Slingshot was used to infer differentiation trajectories (45). Capybara was used to perform quadratic programming as previously described (25,46). Myeloid identity scores were calculated based on differential expression between HSC/MPP and GMP clusters in control sample groups.

### scATAC-seq library construction, sequencing, and analysis

Libraries were generated using Chromium Next GEM Single Cell ATAC Reagent Kits per manufacture instructions and sequenced on an Illumina Novaseq X plus. The Cell Ranger ATAC v2.1.0 pipeline was used to align raw sequencing reads and generate BAM and fragment files. ArchR was then used for downstream scATAC-seq analysis, including quality control, data normalization and dimension reduction, integration with CITE-seq datasets by transferring CITE-seq cell type identities to scATAC-seq cells, cell type-specific peak calling, and ChromVar motif enrichment analysis (47,48).

### EdU incorporation assays

EdU (Invitrogen) was administered by IP injections (100 mg/kg/dose) given every 8 hours beginning 24 hours prior to bone marrow or fetal liver harvest. Assays were performed using the AF488 Click-it Plus EdU Flow Cytometry Assay Kit (Molecular probes, C10633).

### Western blotting

LK cells were sorted into PBS + 0.1% BSA, pelleted, resuspended and transferred to 10% trichloroacetic acid to precipitate protein. Western blots were performed as previously described (49). Anti-FLAG M2 primary antibody (Sigma, F1804) was used to detect FLAG-tagged MLL::ENL from iPSC-derived chimeric LK cells. alpha-Tubulin (Cell Signaling, 2144S) was used as loading control.

### CRISPR/Cas9 editing and transplantations

GMPs were isolated from Tet-OFF-ME mice and cultured overnight in StemSpan SFEM II (Stemcell Technologies) + 10ng/mL murine SCF (PeproTech) + 10ng/mL murine TPO (PeproTech). Individual electroporation reactions were performed for each recipient mouse. Ribonucleoprotein (RNP) complex was prepared by incubating Cas9 (Integrated DNA Technologies) with either *Mll3* gRNA (GTCACTCCAAAAATTGGCATNGG) or a control gRNA (GACATTTCTTTCCCCACTGG) at a 6:1 molar ratio at room temperature for 10 minutes. 4,000 cells were electroporated with RNP and cultured for 6 hours. Electroporated cells were transplanted with 500,000 CD45.1^+^ cells into each lethally irradiated recipient mouse.

### Statistics

Group sizes and statistical tests are cited in each figure legend. The log rank test was used to compare survival curves. Wilcoxon signed-rank test was used for non-parametric comparisons. Other comparisons were made using the Student’s t-test or one-way ANOVA with Tukey’s posthoc test, as indicated. Single cell datasets were analyzed as described in the respective methods sections and figure legends.

### Data availability

All CITE-seq, scATAC-seq and RNA-seq data have been deposited in Gene Expression Omnibus (GSE291041).

## AUTHOR CONTRIBUTIONS

J.A.M. designed and oversaw all experiments, interpreted the data and wrote the manuscript with J.M.-C. J.M.-C. designed, conducted, and interpreted the experiments. W.Y. performed all bioinformatic analyses. E.D., H.C.W., E.B.C., R.M., R.M.P., J.Y., Y.L., and R.C. performed the experiments. L.F.Z.B assisted in establishing the iPSC model. J.M.W. performed blastocyst injections to generate iPSC-derived chimeric mice. All authors reviewed the manuscript.

## ACKOWLEDGEMENTS

We thank David Bryder for providing TetO-MLL::ENL mice. This work was supported by grants to J.A.M. from the National Heart, Lung, and Blood Institute (R01HL152180), the National Cancer Institute (R01CA285272), Gabriel’s Angel Foundation, the Edward P. Evans Foundation and the Children’s Discovery Institute of Washington University and St. Louis Children’s Hospital. J.M.-C. was supported by a National Cancer Institute career development grant (F31CA268923). J.A.M. is a Leukemia and Lymphoma Society Scholar.

## REFERENCES

1. Louka E, Povinelli B, Rodriguez-Meira A, Buck G, Wen WX, Wang G, et al. Heterogeneous disease-propagating stem cells in juvenile myelomonocytic leukemia. J Exp Med 2021;218(2) doi 10.1084/jem.20180853.

2. Biondi A, Cimino G, Pieters R, Pui CH. Biological and therapeutic aspects of infant leukemia. Blood 2000;96(1):24–33.

3. Roberts KG. Genetics and prognosis of ALL in children vs adults. Hematology Am Soc Hematol Educ Program 2018;2018(1):137–45 doi 10.1182/asheducation-2018.1.137.

4. Downing JR, Shannon KM. Acute leukemia: a pediatric perspective. Cancer Cell 2002;2(6):437–45 doi S1535610802002118 [pii].

5. Bolouri H, Farrar JE, Triche T, Jr., Ries RE, Lim EL, Alonzo TA, et al. The molecular landscape of pediatric acute myeloid leukemia reveals recurrent structural alterations and age-specific mutational interactions. Nat Med 2018;24(1):103–12 doi 10.1038/nm.4439.

6. Meyer C, Larghero P, Almeida Lopes B, Burmeister T, Groger D, Sutton R, et al. The KMT2A recombinome of acute leukemias in 2023. Leukemia 2023;37(5):988–1005 doi 10.1038/s41375-023-01877-1.

7. Bernt KM, Zhu N, Sinha AU, Vempati S, Faber J, Krivtsov AV, et al. MLL-rearranged leukemia is dependent on aberrant H3K79 methylation by DOT1L. Cancer Cell 2011;20(1):66–78 doi 10.1016/j.ccr.2011.06.010.

8. Zhu L, Li Q, Wong SH, Huang M, Klein BJ, Shen J, et al. ASH1L Links Histone H3 Lysine 36 Dimethylation to MLL Leukemia. Cancer Discov 2016;6(7):770–83 doi 10.1158/2159-8290.CD-16-0058.

9. Garcia-Cuellar MP, Buttner C, Bartenhagen C, Dugas M, Slany RK. Leukemogenic MLL-ENL Fusions Induce Alternative Chromatin States to Drive a Functionally Dichotomous Group of Target Genes. Cell Rep 2016;15(2):310–22 doi 10.1016/j.celrep.2016.03.018.

10. Umeda M, Ma J, Westover T, Ni Y, Song G, Maciaszek JL, et al. A new genomic framework to categorize pediatric acute myeloid leukemia. Nat Genet 2024;56(2):281–93 doi 10.1038/s41588-023-01640-3.

11. Andersson AK, Ma J, Wang J, Chen X, Gedman AL, Dang J, et al. The landscape of somatic mutations in infant MLL-rearranged acute lymphoblastic leukemias. Nat Genet 2015;47(4):330–7 doi 10.1038/ng.3230.

12. Greaves MF, Wiemels J. Origins of chromosome translocations in childhood leukaemia. Nat Rev Cancer 2003;3(9):639–49 doi 10.1038/nrc1164.

13. Wiemels JL, Cazzaniga G, Daniotti M, Eden OB, Addison GM, Masera G, et al. Prenatal origin of acute lymphoblastic leukaemia in children. Lancet 1999;354(9189):1499–503 doi S0140673699094039 [pii].

14. Gale KB, Ford AM, Repp R, Borkhardt A, Keller C, Eden OB, et al. Backtracking leukemia to birth: identification of clonotypic gene fusion sequences in neonatal blood spots. Proc Natl Acad Sci U S A 1997;94(25):13950–4.

15. Ford AM, Ridge SA, Cabrera ME, Mahmoud H, Steel CM, Chan LC, et al. In utero rearrangements in the trithorax-related oncogene in infant leukaemias. Nature 1993;363(6427):358–60 doi 10.1038/363358a0.

16. Gill Super HJ, Rothberg PG, Kobayashi H, Freeman AI, Diaz MO, Rowley JD. Clonal, nonconstitutional rearrangements of the MLL gene in infant twins with acute lymphoblastic leukemia: in utero chromosome rearrangement of 11q23. Blood 1994;83(3):641–4.

17. Campbell M, Cabrera ME, Legues ME, Ridge S, Greaves M. Discordant clinical presentation and outcome in infant twins sharing a common clonal leukaemia. Br J Haematol 1996;93(1):166–9 doi 10.1046/j.1365-2141.1996.455999.x.

18. Horton SJ, Jaques J, Woolthuis C, van Dijk J, Mesuraca M, Huls G, et al. MLL-AF9-mediated immortalization of human hematopoietic cells along different lineages changes during ontogeny. Leukemia 2013;27(5):1116–26 doi 10.1038/leu.2012.343.

19. Secker KA, Bruns L, Keppeler H, Jeong J, Hentrich T, Schulze-Hentrich JM, et al. Only Hematopoietic Stem and Progenitor Cells from Cord Blood Are Susceptible to Malignant Transformation by MLL-AF4 Translocations. Cancers (Basel) 2020;12(6) doi 10.3390/cancers12061487.

20. Okeyo-Owuor T, Li Y, Patel RM, Yang W, Casey EB, Cluster AS, et al. The efficiency of murine MLL-ENL-driven leukemia initiation changes with age and peaks during neonatal development. Blood Adv 2019;3(15):2388–99 doi 10.1182/bloodadvances.2019000554.

21. Islami F, Ward EM, Sung H, Cronin KA, Tangka FKL, Sherman RL, et al. Annual Report to the Nation on the Status of Cancer, Part 1: National Cancer Statistics. J Natl Cancer Inst 2021 doi 10.1093/jnci/djab131.

22. Isaacs H, Jr. Fetal and neonatal leukemia. J Pediatr Hematol Oncol 2003;25(5):348–61 doi 10.1097/00043426-200305000-00002.

23. Eldeeb M, Yuan O, Guzzi N, Thi Ngoc PC, Konturek-Ciesla A, Kristiansen TA, et al. A fetal tumor suppressor axis abrogates MLL-fusion-driven acute myeloid leukemia. Cell Rep 2023;42(2):112099 doi 10.1016/j.celrep.2023.112099.

24. Li Y, Mendoza-Castrejon J, Patel RM, Casey EB, Denby E, Bryder D, et al. LIN28B promotes differentiation of fully transformed AML cells but is dispensable for fetal leukemia suppression. Leukemia 2024;38(3):648–51 doi 10.1038/s41375-024-02167-0.

25. Kong W, Fu YC, Holloway EM, Garipler G, Yang X, Mazzoni EO, et al. Capybara: A computational tool to measure cell identity and fate transitions. Cell Stem Cell 2022;29(4):635–49 e11 doi 10.1016/j.stem.2022.03.001.

26. Zhang H, Alberich-Jorda M, Amabile G, Yang H, Staber PB, Diruscio A, et al. Sox4 is a key oncogenic target in C/EBPalpha mutant acute myeloid leukemia. Cancer Cell 2013;24(5):575–88 doi 10.1016/j.ccr.2013.09.018.

27. Brown FC, Still E, Koche RP, Yim CY, Takao S, Cifani P, et al. MEF2C Phosphorylation Is Required for Chemotherapy Resistance in Acute Myeloid Leukemia. Cancer Discov 2018;8(4):478–97 doi 10.1158/2159-8290.CD-17-1271.

28. Baker SJ, Ma’ayan A, Lieu YK, John P, Reddy MV, Chen EY, et al. B-myb is an essential regulator of hematopoietic stem cell and myeloid progenitor cell development. Proc Natl Acad Sci U S A 2014;111(8):3122–7 doi 10.1073/pnas.1315464111.

29. Cheng JC, Kinjo K, Judelson DR, Chang J, Wu WS, Schmid I, et al. CREB is a critical regulator of normal hematopoiesis and leukemogenesis. Blood 2008;111(3):1182–92 doi 10.1182/blood-2007-04-083600.

30. Almotiri A, Alzahrani H, Menendez-Gonzalez JB, Abdelfattah A, Alotaibi B, Saleh L, et al. Zeb1 modulates hematopoietic stem cell fates required for suppressing acute myeloid leukemia. J Clin Invest 2021;131(1) doi 10.1172/JCI129115.

31. Chen R, Okeyo-Owuor T, Patel RM, Casey EB, Cluster AS, Yang W, et al. Kmt2c mutations enhance HSC self-renewal capacity and convey a selective advantage after chemotherapy. Cell Rep 2021;34(7):108751 doi 10.1016/j.celrep.2021.108751.

32. Wang HC, Chen R, Yang W, Li Y, Muthukumar R, Patel RM, et al. Kmt2c restricts G-CSF-driven HSC mobilization and granulocyte production in a methyltransferase-independent manner. Cell Rep 2024;43(8):114542 doi 10.1016/j.celrep.2024.114542.

33. Soto-Feliciano YM, Sanchez-Rivera FJ, Perner F, Barrows DW, Kastenhuber ER, Ho YJ, et al. A Molecular Switch between Mammalian MLL Complexes Dictates Response to Menin-MLL Inhibition. Cancer Discov 2023;13(1):146–69 doi 10.1158/2159-8290.CD-22-0416.

34. Dorighi KM, Swigut T, Henriques T, Bhanu NV, Scruggs BS, Nady N, et al. Mll3 and Mll4 Facilitate Enhancer RNA Synthesis and Transcription from Promoters Independently of H3K4 Monomethylation. Mol Cell 2017;66(4):568–76 e4 doi 10.1016/j.molcel.2017.04.018.

35. Ugale A, Norddahl GL, Wahlestedt M, Sawen P, Jaako P, Pronk CJ, et al. Hematopoietic stem cells are intrinsically protected against MLL-ENL-mediated transformation. Cell Rep 2014;9(4):1246–55 doi 10.1016/j.celrep.2014.10.036.

36. Vogelstein B, Kinzler KW. The multistep nature of cancer. Trends Genet 1993;9(4):138–41 doi 10.1016/0168-9525(93)90209-z.

37. Li Y, Yang W, Patel RM, Casey EB, Denby E, Mendoza-Castrejon J, et al. FLT3ITD drives context-specific changes in cell identity and variable interferon dependence during AML initiation. Blood 2023;141(12):1442–56 doi 10.1182/blood.2022016889.

38. Li Y, Yang W, Wang HC, Patel RM, Casey EB, Denby E, et al. Basal type I interferon signaling has only modest effects on neonatal and juvenile hematopoiesis. Blood Adv 2023;7(11):2609–21 doi 10.1182/bloodadvances.2022008595.

39. Venkatasubramanian M, Chetal K, Schnell DJ, Atluri G, Salomonis N. Resolving single-cell heterogeneity from hundreds of thousands of cells through sequential hybrid clustering and NMF. Bioinformatics 2020;36(12):3773–80 doi 10.1093/bioinformatics/btaa201.

40. Kang JB, Nathan A, Weinand K, Zhang F, Millard N, Rumker L, et al. Efficient and precise single-cell reference atlas mapping with Symphony. Nat Commun 2021;12(1):5890 doi 10.1038/s41467-021-25957-x.

41. Korsunsky I, Nathan A, Millard N, Raychaudhuri S. Presto scales Wilcoxon and auROC analyses to millions of observations. bioRxiv 2019 doi 10.1101/653253.

42. Love MI, Huber W, Anders S. Moderated estimation of fold change and dispersion for RNA-seq data with DESeq2. Genome Biol 2014;15(12):550 doi 10.1186/s13059-014-0550-8.

43. Subramanian A, Tamayo P, Mootha VK, Mukherjee S, Ebert BL, Gillette MA, et al. Gene set enrichment analysis: a knowledge-based approach for interpreting genome-wide expression profiles. Proc Natl Acad Sci U S A 2005;102(43):15545–50 doi 0506580102 [pii] 10.1073/pnas.0506580102.

44. Hanzelmann S, Castelo R, Guinney J. GSVA: gene set variation analysis for microarray and RNA-seq data. BMC Bioinformatics 2013;14:7 doi 10.1186/1471-2105-14-7.

45. Street K, Risso D, Fletcher RB, Das D, Ngai J, Yosef N, et al. Slingshot: cell lineage and pseudotime inference for single-cell transcriptomics. BMC Genomics 2018;19(1):477 doi 10.1186/s12864-018-4772-0.

46. Li Y, Kong W, Yang W, Patel RM, Casey EB, Okeyo-Owuor T, et al. Single-Cell Analysis of Neonatal HSC Ontogeny Reveals Gradual and Uncoordinated Transcriptional Reprogramming that Begins before Birth. Cell Stem Cell 2020;27(5):732–47 doi 10.1016/j.stem.2020.08.001.

47. Granja JM, Corces MR, Pierce SE, Bagdatli ST, Choudhry H, Chang HY, et al. ArchR is a scalable software package for integrative single-cell chromatin accessibility analysis. Nat Genet 2021;53(3):403–11 doi 10.1038/s41588-021-00790-6.

48. Schep AN, Wu B, Buenrostro JD, Greenleaf WJ. chromVAR: inferring transcription-factor-associated accessibility from single-cell epigenomic data. Nat Methods 2017;14(10):975–8 doi 10.1038/nmeth.4401.

49. Porter SN, Cluster AS, Yang W, Busken KA, Patel RM, Ryoo J, et al. Fetal and neonatal hematopoietic progenitors are functionally and transcriptionally resistant to Flt3-ITD mutations. Elife 2016;5:e18882 doi 10.7554/eLife.18882.

