## Supplemental materials for "Fetal context conveys heritable protection against MLL-rearranged leukemia that depends on MLL3"

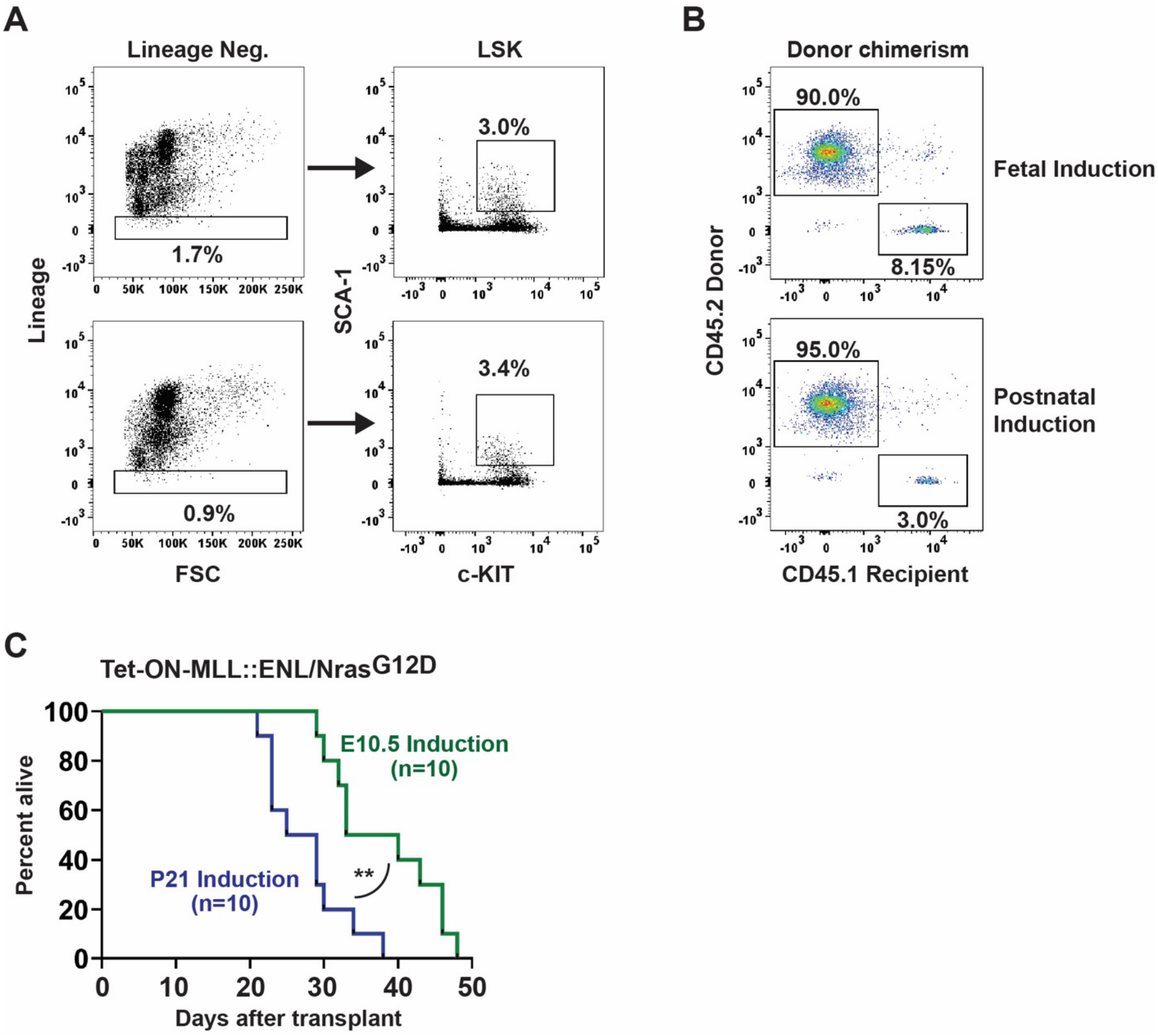


**Supplemental Figure 1 (related to Figure 1). Tet-On-ME/Nras$^{G12D}$ transplant and flow cytometry gating for MLL-ENL-expressing progenitors.** (A) Flow cytometry gating strategy for sorting donor LSK cells. (B) Flow cytometry plots from peripheral blood of recipient mice (Figure 1D) showing donor (CD45.2) chimerism. (C) Kaplan-Meier survival curves for recipients of Tet-On-ME; Nras$^{G12D}$ LSK cells after MLL::ENL induction at E10.5 or P21 and transplantation at P28. n=10 per group, **$p<0.01$ by log rank test.

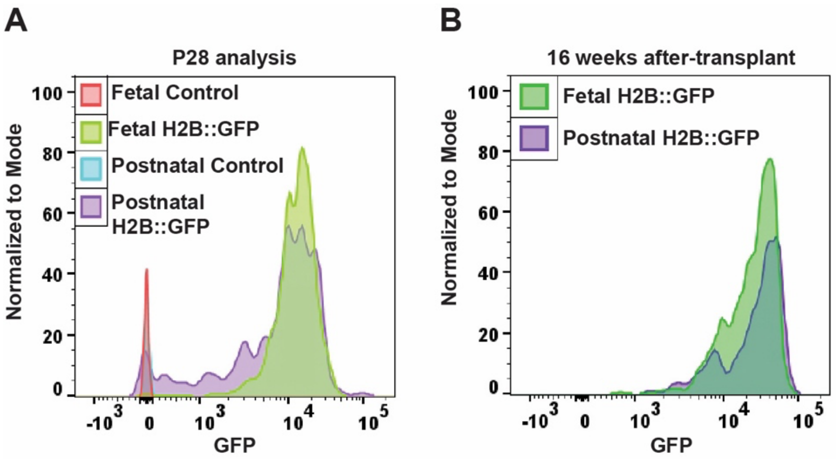


**Supplemental Figure 2 (related to Figure 1). Selection against MLL::ENL transgene expression is not observed for other transgenes integrated within the *Col1a1* locus.** (A) H2B::GFP expression in P28 Tet-Off-H2B-GFP (Vav1-Cre; Rosa26$^{LoxP\text{-}STOP\text{-}LoxP_tTA}$; Col1a1$^{TetO_H2B::GFP}$) mouse bone marrow after fetal or postnatal H2B-GFP induction. In contrast to the Tet-Off-MLL::ENL model, H2B-GFP expression levels were comparable at P28 after fetal and postnatal induction. (B) H2B-GFP expression in recipient mouse bone marrow 16 weeks after transplantation. Donor Tet-Off-H2B-GFP mice had H2B-GFP induced either during fetal or postnatal stages. In contrast to the Tet-Off-MLL::ENL model, H2B-GFP expression levels were comparable between the groups.

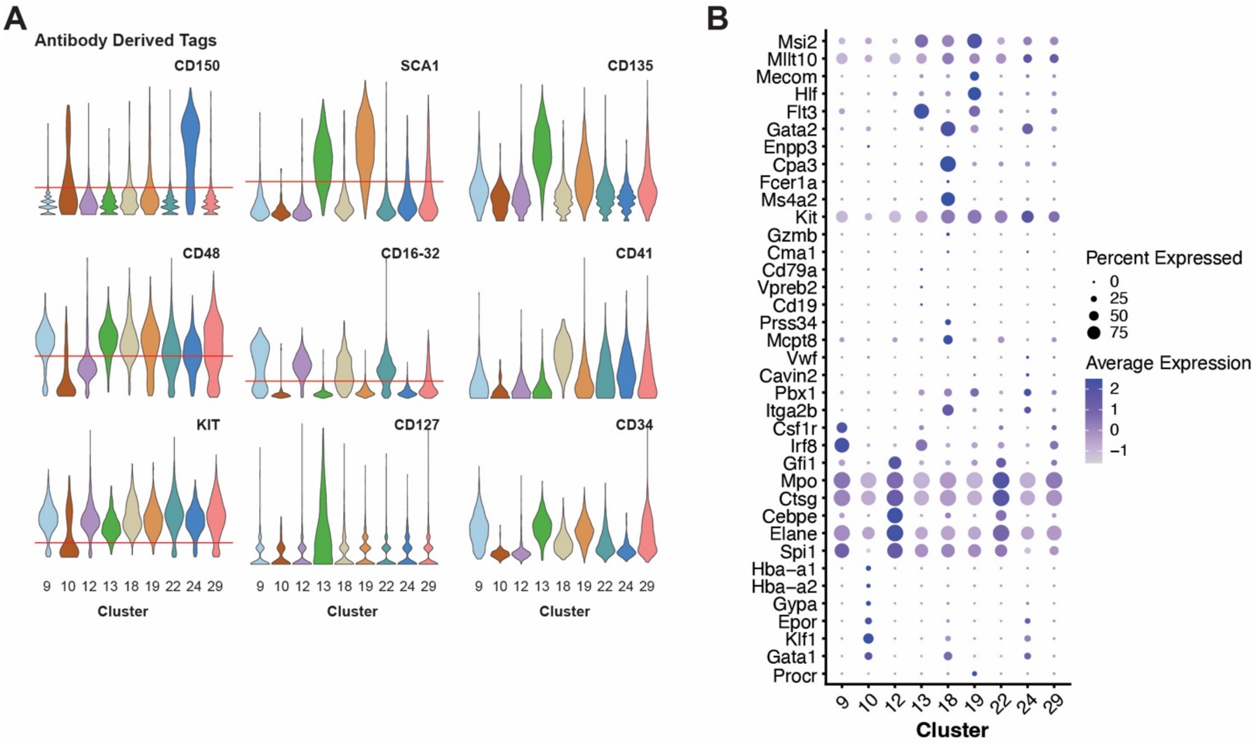


**Supplemental Figure 3 (related to Figure 2). Annotation strategies for P28 CITE-seq data.** (A) Expression of indicated surface markers based on Antibody Derived Tags. Expression levels are shown for each cluster in Figure 2A. CD150, CD48, SCA1, CD16-31 and KIT were used to annotate HSC/MPP and pGM/GMP clusters. Thresholds for positive versus negative expression are shown in red. (B) For some clusters, marker gene expression was used to inform cluster annotation. Marker expression is shown.

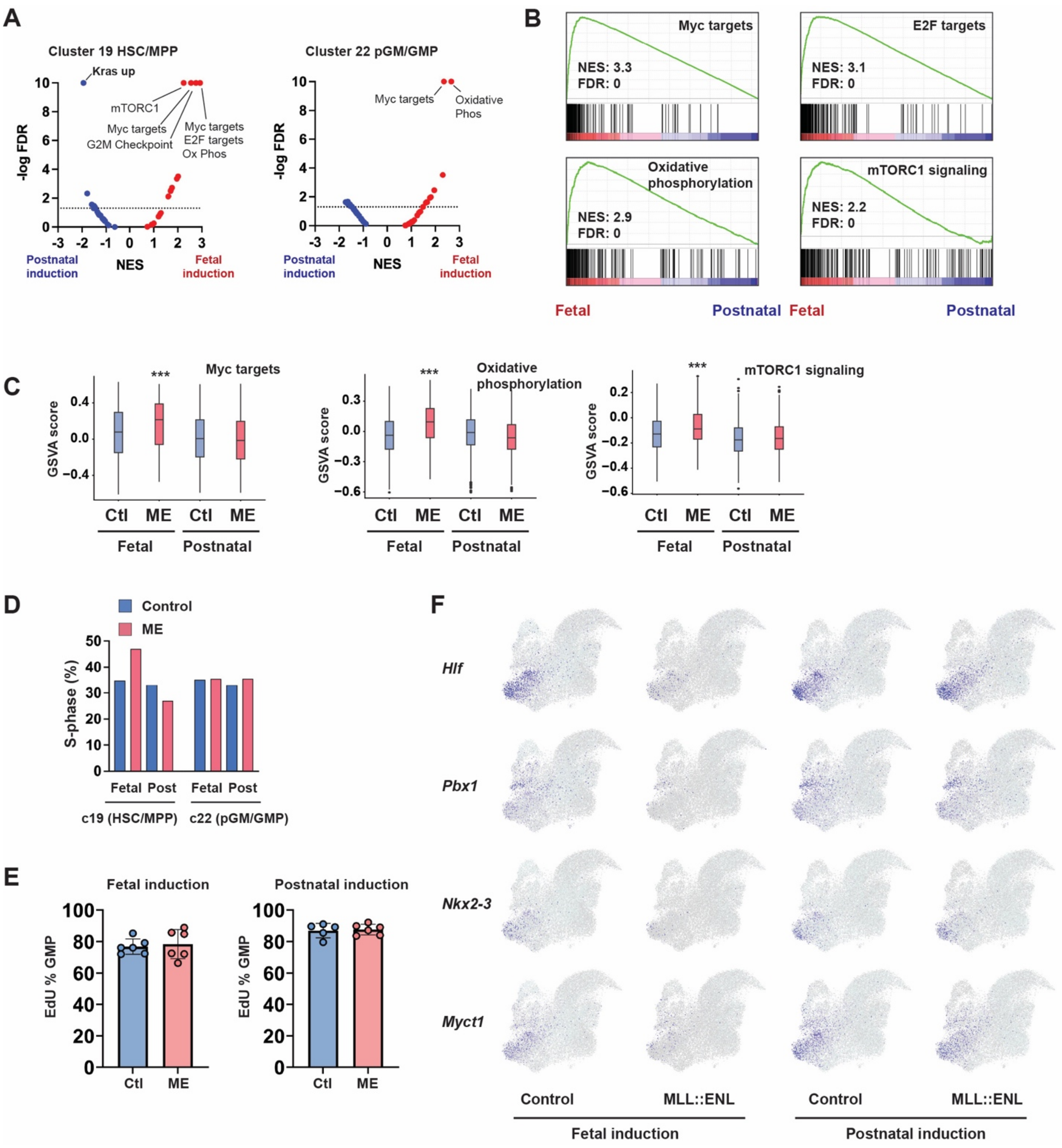


**Supplemental Figure 4 (related to Figure 2). Fetal MLL::ENL induction leads to enhanced protein synthesis and oxidative metabolism in postnatal MPPs while dampening self-renewal gene expression.** (A) Volcano plots showing normalized enrichment scores and false discovery rates for Hallmark gene sets, as calculated from 4 pseudoreplicates per genotype for the HSC/MPP and pGM/GMP populations. (B) GSEA plots for Hallmark Myc targets, oxidative phosphorylation, E2F targets and mTORC1 signaling genes sets, comparing MLL::ENL-expressing (ME) HSC/MPPs after fetal or postnatal induction. (C) Box plots showing gene set variance analysis (GSVA) scores for control and MLL::ENL-expressing HSC/MPPs after fetal or postnatal induction. (D) Percent of cells in S-phase based on gene expression in HSC/MPP and pGM/GMP clusters for indicated genotype and age of MLL::ENL induction. (E) Percent of EdU+ GMPs in P28 control and MLL::ENL-expressing mice after fetal or postnatal induction. (F) UMAPs showing single cell expression of *Hlf*, *Pbx1*, *Nkx2-3*, and *Myct1* genes for indicated genotypes and ages of MLL::ENL induction.

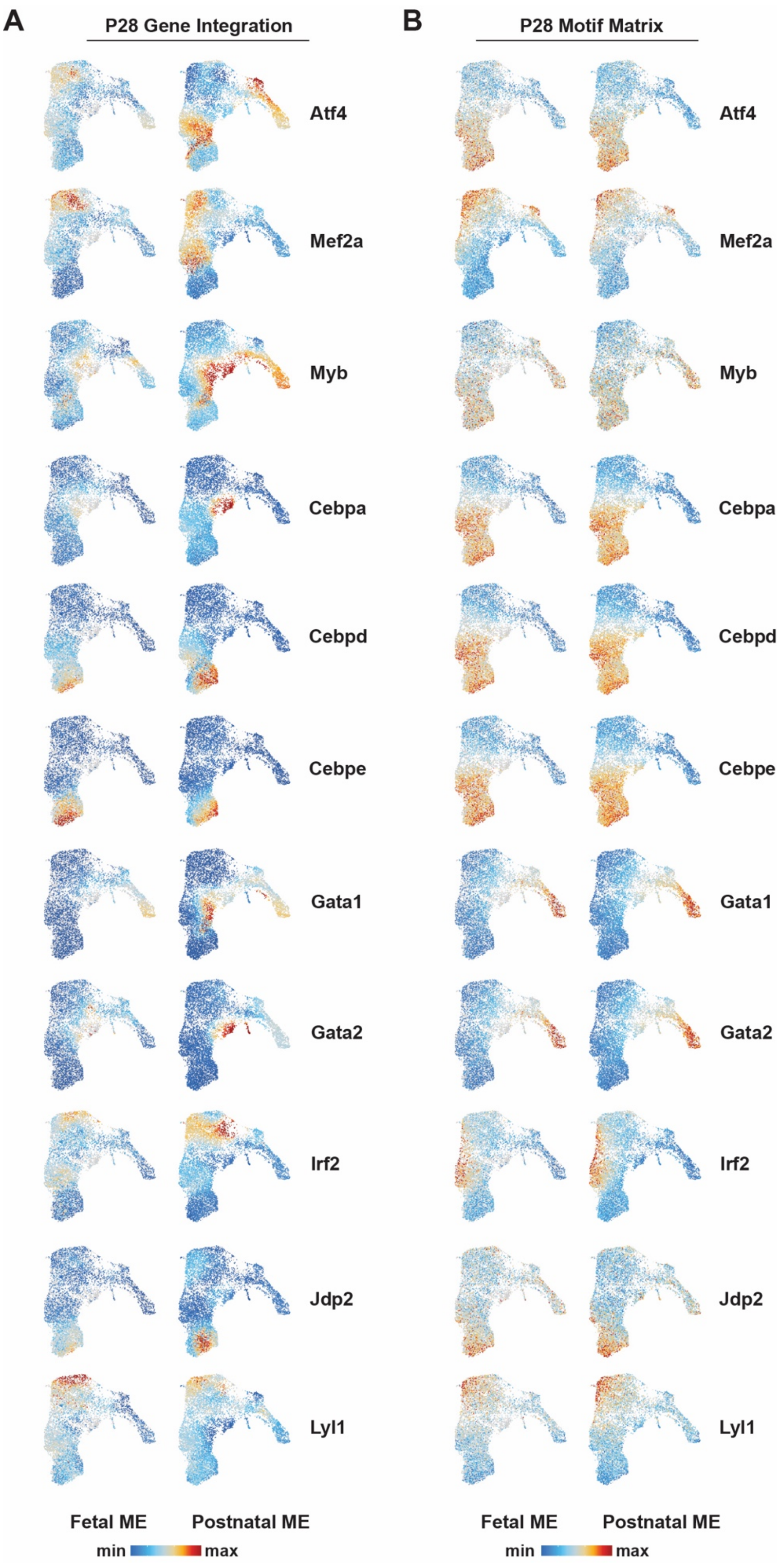
A
P28 Gene Integration
B
P28 Motif Matrix
Atf4
Mef2a
Myb
Cebpa
Cebpd
Cebpe
Gata1
Gata2
Irf2
Jdp2
Lyl1
Fetal ME
Postnatal ME
min
max

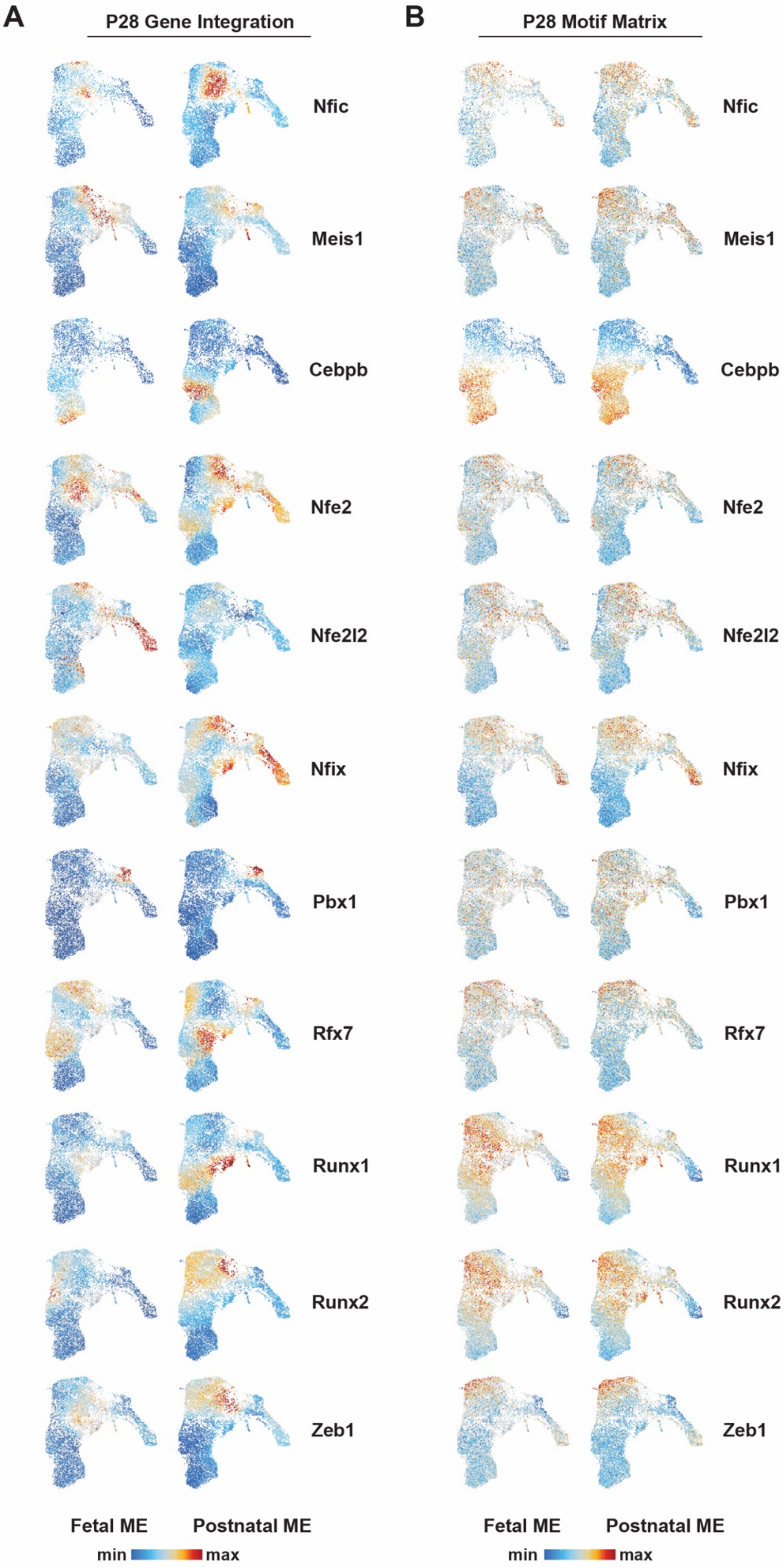
A
P28 Gene Integration
B
P28 Motif Matrix
Nfic
Meis1
Cebpb
Nfe2
Nfe2l2
Nfix
Pbx1
Rfx7
Runx1
Runx2
Zeb1
Fetal ME
Postnatal ME
min
max

**Supplemental Figure 5 (related to Figure 3). MLL::ENL induction leads to changes in gene expression at P28 without corresponding changes in chromatin accessibility.** (A) Transcript expression for indicated genes in individual cells, projected as heatmaps on the UMAPs from panel A. Ranges of min/max expression are identical for fetal and postnatal induction cohorts for each gene. (B) Motif enrichment for indicated transcription factors within individual cells, projected as heatmaps. Ranges of min/max expression are identical for fetal and postnatal induction cohorts for each gene.

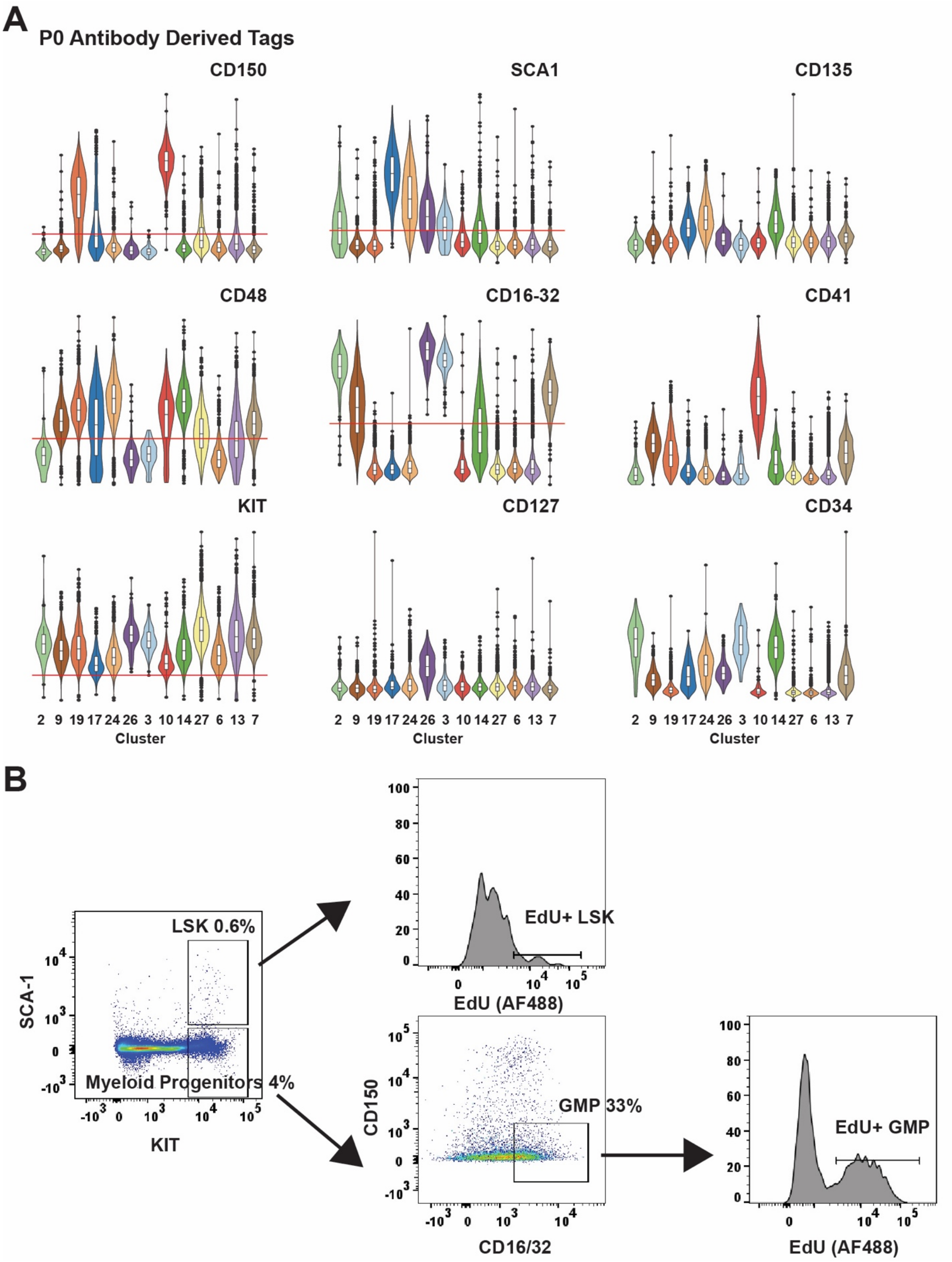
A
P0 Antibody Derived Tags
CD150
SCA1
CD135
CD48
CD16-32
CD41
KIT
CD127
CD34
2 9 19 17 24 26 3 10 14 27 6 13 7
Cluster
B
LSK 0.6%
Myeloid Progenitors 4%
SCA-1
KIT
EdU+ LSK
EdU (AF488)
GMP 33%
CD150
CD16/32
EdU+ GMP
EdU (AF488)

**Supplemental Figure 6 (related to Figure 4). Annotation of P0 CITE-seq clusters and gating strategy for EdU assays.** (A) Expression of indicated surface markers based on ADT. Expression levels are shown for each cluster in Figure 4A. CD150, CD48, SCA1, CD16-31 and KIT were used to annotate HSC/MPP and pGM/GMP clusters. Thresholds for positive versus negative expression are shown in red. (B) Flow cytometry gating strategy for assessing EdU+ LSK and EdU+ GMP.

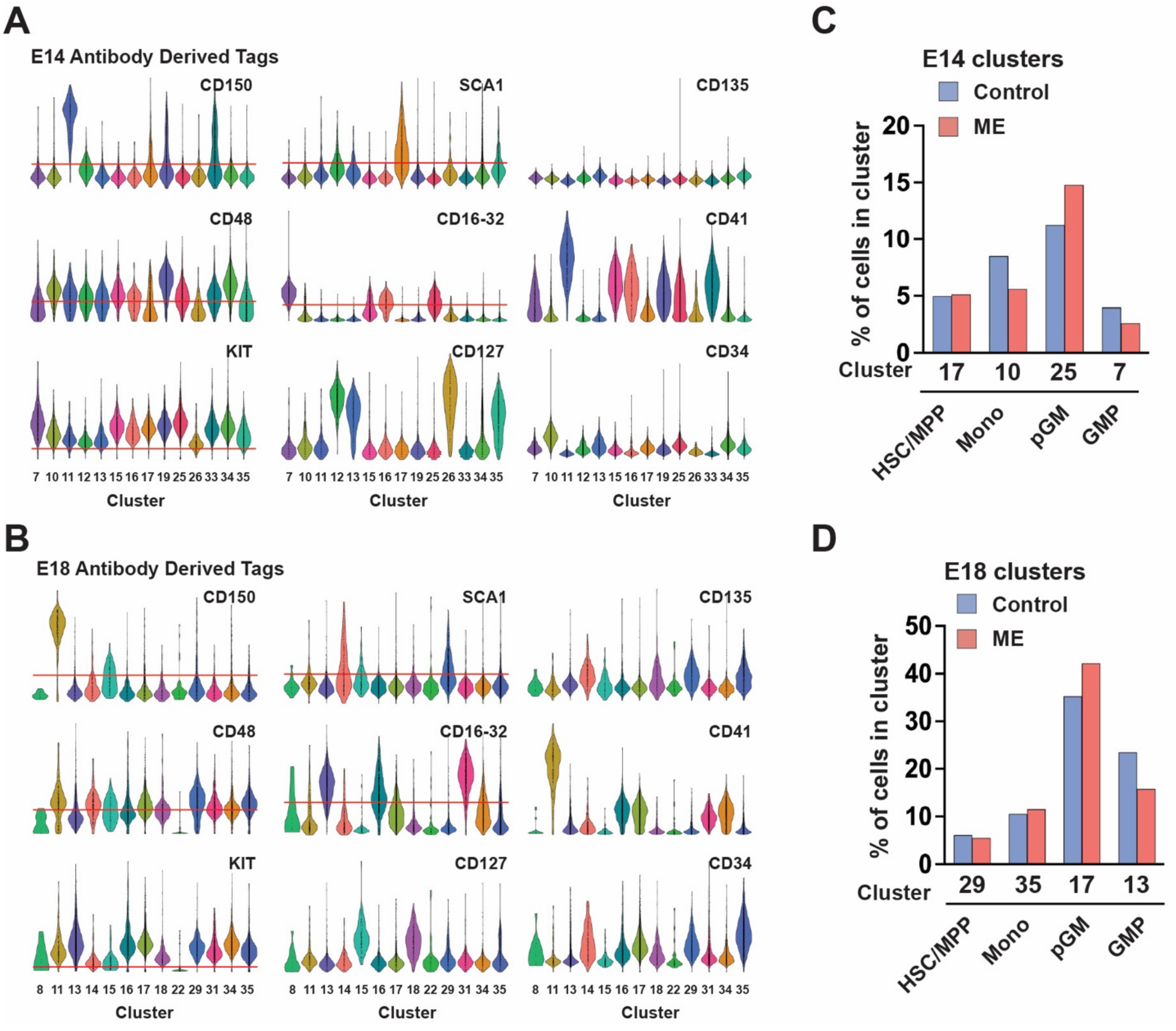


**Supplemental Figure 7 (related to Figure 5). Annotation of fetal CITE-seq clusters and cluster distributions.** (A, B) Expression of indicated surface markers based on ADT. Expression levels are shown for each cluster in Figure 5D, E. CD150, CD48, SCA1, CD16-31 and KIT were used to annotate HSC/MPP and pGM/GMP clusters. Thresholds for positive versus negative expression are shown in red. (C, D) Distribution of cells within indicated clusters at E14 (C) and E18 (D).

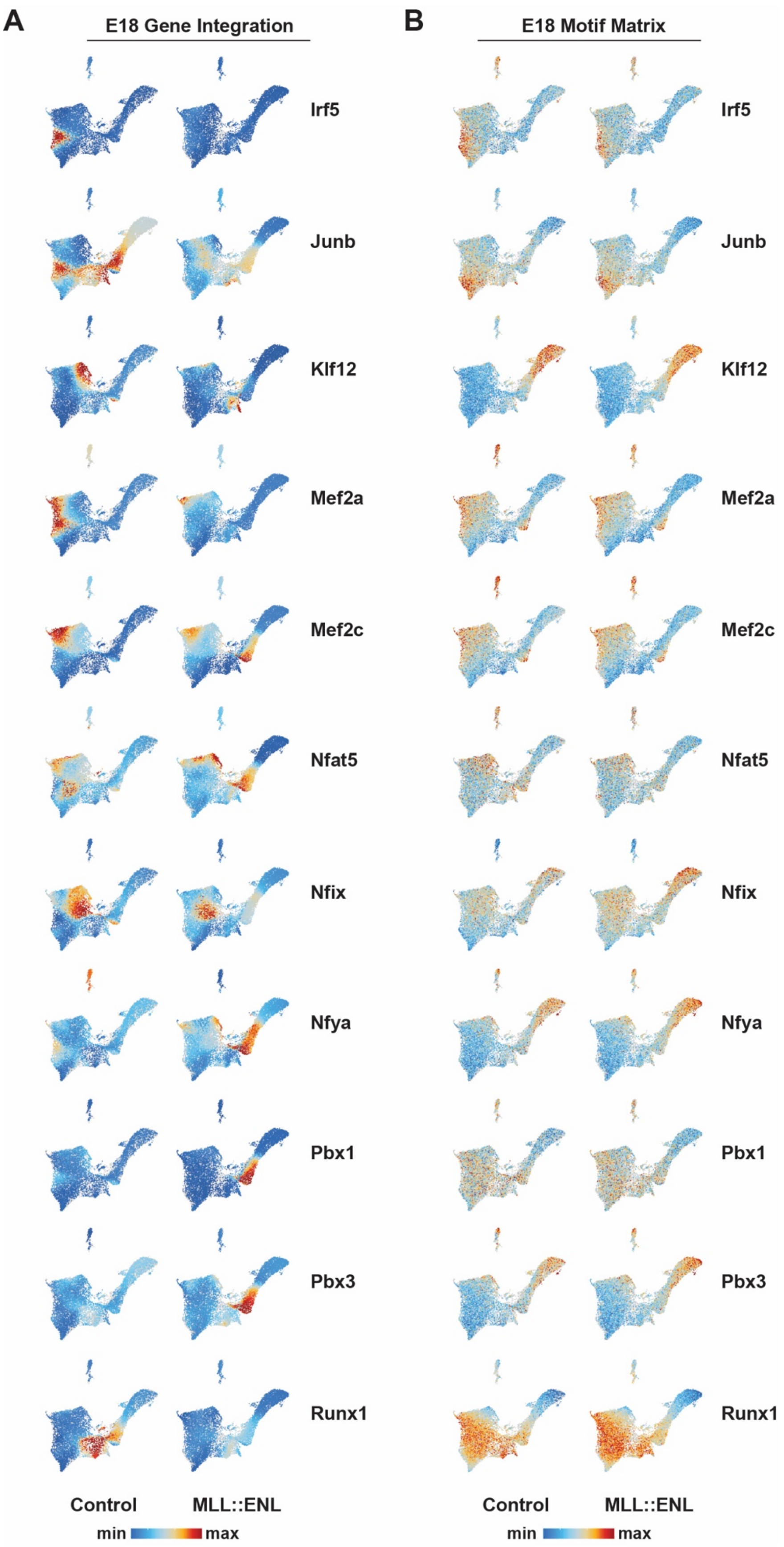
A
E18 Gene Integration
B
E18 Motif Matrix
Irf5
Junb
Klf12
Mef2a
Mef2c
Nfat5
Nfix
Nfya
Pbx1
Pbx3
Runx1
Control
MLL::ENL
min
max

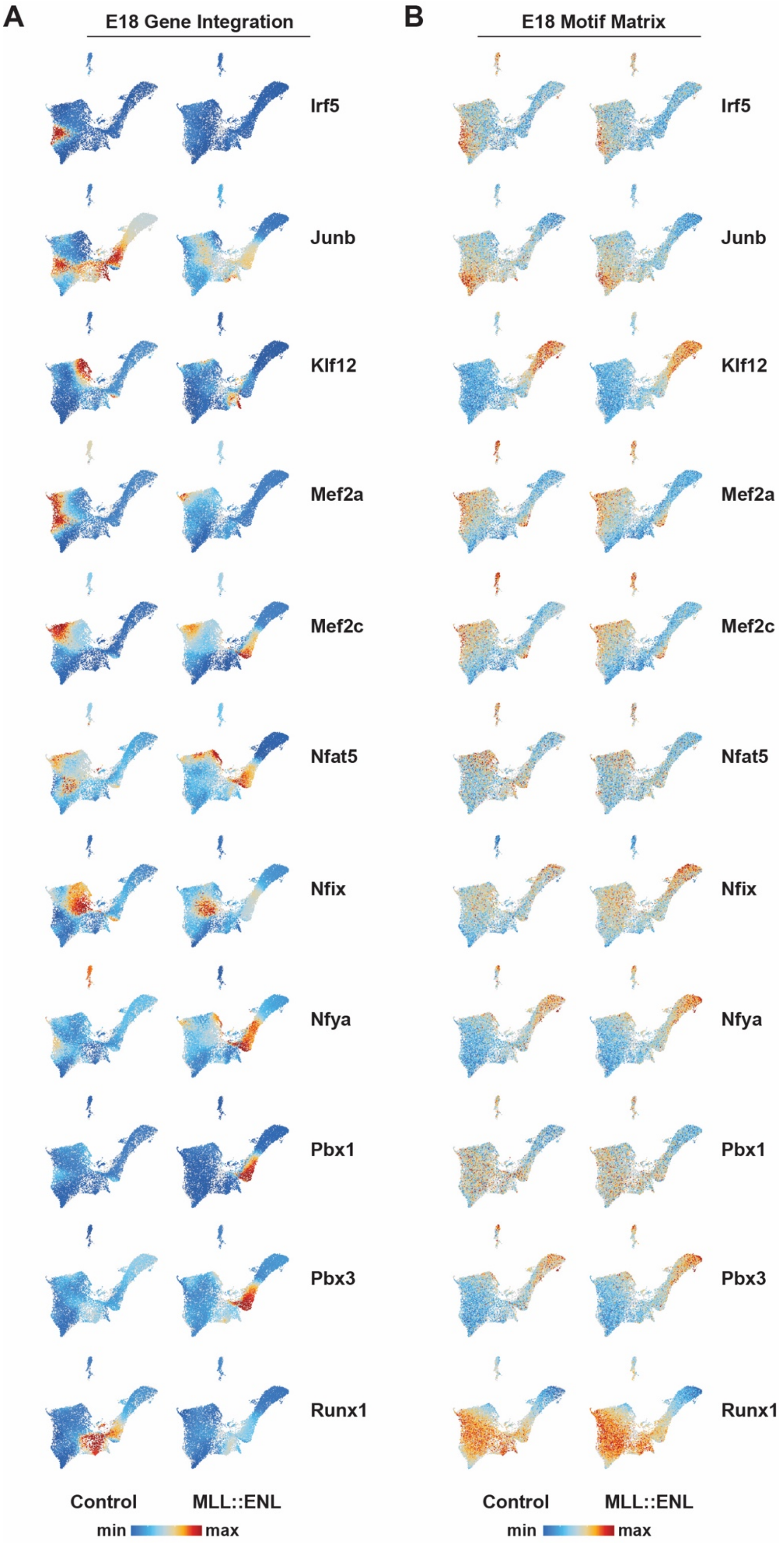
A
E18 Gene Integration
B
E18 Motif Matrix
Irf5
Junb
Klf12
Mef2a
Mef2c
Nfat5
Nfix
Nfya
Pbx1
Pbx3
Runx1
Control
MLL::ENL
min
max

**Supplemental Figure 8 (related to Figure 6).** (A) Transcript expression for indicated genes in individual cells, projected as heatmaps on the UMAPs, for E18 cells. Ranges of min/max expression are identical for control and MLL::ENL-expressing cohorts for each gene. (B) Motif enrichment for indicated transcription factors within individual cells, projected as heatmaps. Ranges of min/max expression are identical for control and MLL::ENL-expressing cohorts for each gene.
